# CoRa –A general approach for quantifying biological feedback control

**DOI:** 10.1101/2020.10.09.334078

**Authors:** Mariana Gómez-Schiavon, Hana El-Samad

**Affiliations:** Department of Biochemistry and Biophysics, University of California, San Francisco, San Francisco, CA, USA; Cell Design Initiative, University of California, San Francisco, CA, USA; Chan–Zuckerberg Biohub, San Francisco, CA, USA; Cell Design Institute, San Francisco, CA, USA; Laboratorio Internacional de Investigación sobre el Genoma Humano, Universidad Nacional Autónoma de México, Santiago de Querétaro, México; ANID—Millennium Science Initiative Program—Millennium Institute for Integrative Biology (iBio), Santiago 8331150, Chile

## Abstract

Feedback control is a fundamental underpinning of life, underlying homeostasis of biological processes at every scale of organization, from cells to ecosystems. The ability to evaluate the contribution and limitations of feedback control mechanisms operating in cells is a critical step for understanding and ultimately designing feedback control systems with biological molecules. Here, we introduce **CoRa** –or ***Co****ntrol* ***Ra****tio*–, a general framework that quantifies the contribution of a biological feedback control mechanism to adaptation using a mathematically controlled comparison to an identical system that does not contain the feedback. **CoRa** provides a simple and intuitive metric with broad applicability to biological feedback systems.

---

Feedback control is a mechanism by which a system can assess its own state and use this information to react accordingly [15]. Cells and organisms make abundant use of feedback control [3], in particular negative feedback to deploy corrective actions. Negative feedback is instrumental in the ability of biological systems to restore homeostasis after a perturbation [5, 8–10, 17], a property known in engineering as disturbance rejection and in the biological sciences as adaptation. Despite the importance of feedback, no systematic and generalizable approaches exist to quantify the contribution of a negative feedback loop to adaptation in biological networks. Here, we propose **CoRa** –or **Co**ntrol **Ra**tio–, a mathematical approach that tackles this problem. CoRa follows the classical notion of Mathematically Controlled Comparisons [1] by assessing the performance of a biological system with feedback control to a locally analogous system without feedback. The *locally analogous* system without feedback has identical structure and parameters to those of the feedback system, except for the feedback link, and both systems rest at the same steady-state value before the perturbation. As a result, the divergence in their behavior after they are challenged with a perturbation isolates and quantifies the contribution of the feedback control (Fig. 1). CoRa can be defined and computed for any biological system described by a solvable set of ordinary differential equations, irrespective of its complexity. CoRa can also be efficiently computed across different parameter values of a system, allowing a global view of the performance of its feedback under different conditions.

**Figure 1.**
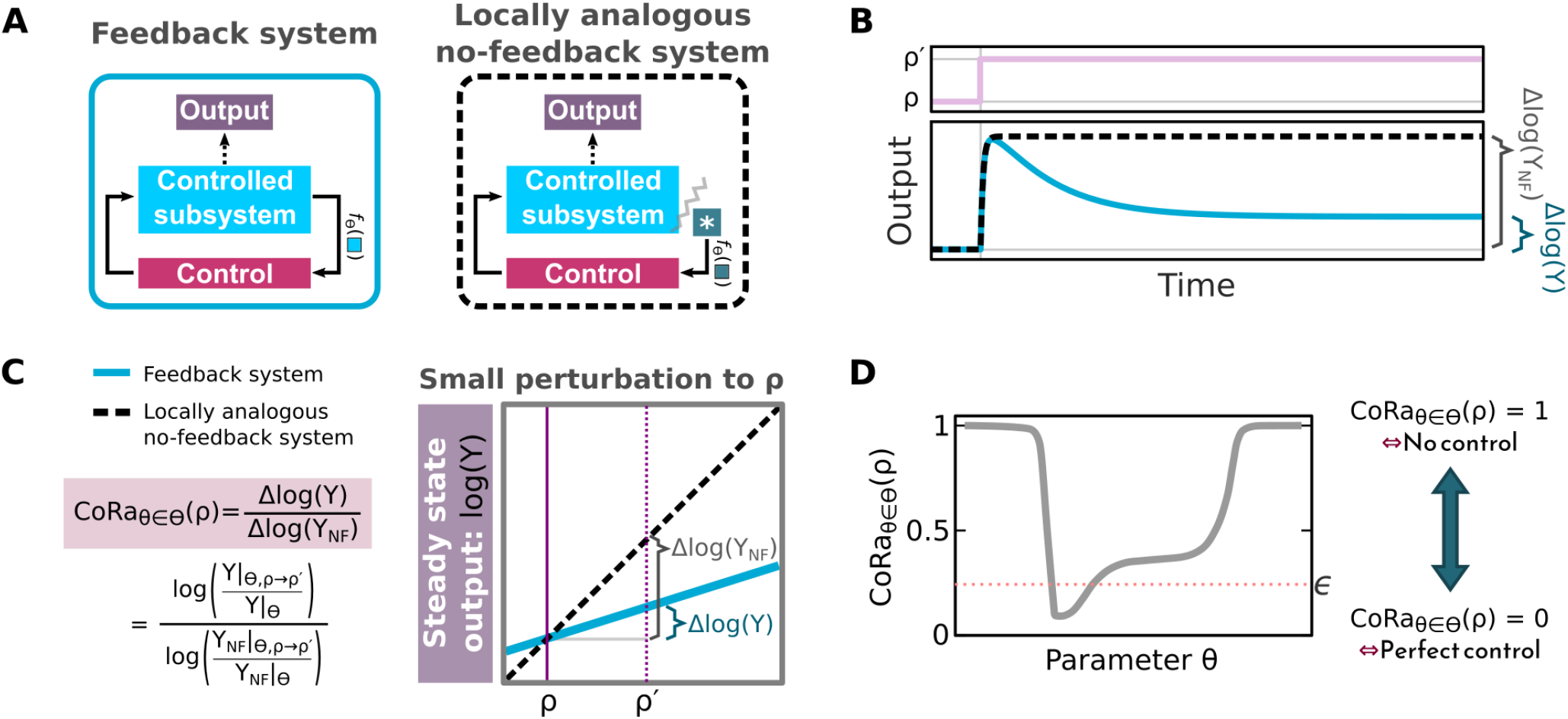
Explaining CoRa, *Control Ratio* approach. **(A)** Diagram of a system with feedback (left) and its locally analogous system without feedback (right). The controlled subsystem is the biological network to which feedback control is applied. For a given parameter set (Θ), the constant input is fixed such that the input signal (*f*_Θ_) is the same between the feedback system and the locally analogous system without feedback. **(B)** Plot of the output of the system with feedback (blue line) and the locally analogous system (black dashed line) as functions of time after a small perturbation to a specific parameter (*ρ* → *ρ*′, with *ρ* ∈ Θ). Since both systems are identical before the perturbation, any differential output response reflects the properties of the feedback. *Y* is used to denote output of system with feedback and *Y*_*NF*_ the output of locally analogous system with no feedback. **(C)** Definition of CoRa. For each parameter set Θ, the CoRa value for perturbation to *ρ*, CoRa_*θ*∈Θ_(*ρ*), is defined as the ratio of the output change of the feedback (Δlog(*Y*)) and no-feedback locally analogous (Δlog(*Y*_*NF*_)) systems after a small perturbation (*ρ* → *ρ* ′). Left panel gives the formula for CoRa and the right panel shows a graphical interpretation of this quantity. (D) CoRa_*θ*∈Θ_(*ρ*) for perturbations in *ρ* can be calculated across a range of values of a chosen parameter *θ* ∈ Θ. This function gives insight as to potency of control for different values of the parameters. For example, one can specify a defined threshold (*ϵ*) for CoRa as an acceptable performance metric, and explore the parameter range for which the control is better than *ϵ*.

## CoRa formalism

To apply CoRa, two systems are considered: the intact system that has the feedback structure, and a locally analogous system without feedback, each of them described by a set of ordinary differential equations with parameters Θ. The locally analogous system is designed to have exactly the same biochemical reactions as the feedback system, with the only difference being the removal of the direct influence of the system output (*Y*) over the rest of the system, therefore generating a system without feedback. Instead, a constant input is introduced in the locally analogous system that mimics the direct influence of *Y* on the relevant chemical species in the system. This positions both systems, the feedback and its locally analogous system with no feedback, at identical steady-state values for the output and all internal variables under the given parameter set Θ. A step-by-step procedure for generating such an analogous system is detailed in *Supplementary Information*.

Once the feedback system and locally analogous system without feedback are defined in this way, the broad idea of CoRa is that in order to evaluate the contribution of the feedback to adaptation following a perturbation in a specific parameter *ρ* ∈ Θ, one can apply a small perturbation (*ρ* → *ρ* ′) and compare the output of the two systems. With *Y* defined as the output of the system with feedback and *Y*_*NF*_ that of the locally analogous system without feedback, the contribution of the feedback to adaptation after a perturbation in parameter *ρ* ∈ Θ can be quantified as the ratio of the response of the feedback system (Δlog(*Y*)) and its locally analogous system (Δlog(*Y*_*NF*_)), 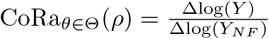 (Fig. 1C, the *θ* ∈ Θ notation is added to indicate that CoRa can be computed for different parameters of the system). Being locally analogous, the two compared systems possess the same nonlinearities and saturations under the given parameter set Θ. As a result, any differences in the response to a small perturbation are attributed to the effect of feedback (*Supplementary Information*).

CoRa provides an easily interpretable assessment of how a system with feedback, positioned at the parameter set Θ, fares compared to a no-feedback system when *ρ* is perturbed. For instance, if CoRa_*θ*∈Θ_(*ρ*) ∈ [0,1), the presence of the feedback reduces the effect of the perturbation compared to the locally analogous system without feedback, Δlog(*Y*) < Δlog(*Y*_*NF*_) (Fig. S1). When CoRa_*θ*∈Θ_(*ρ*) = 0, the feedback endows the system with perfect adaptation (Δlog(*Y*) = 0), with the output returning exactly to the pre-perturbed state even in the continued presence of the perturbation. The value of CoRa_*θ*∈Θ_(*ρ*) increases as the control effect decreases, and when CoRa_*θ*∈Θ_(*ρ*) = 1, the feedback is ineffective as the output of the system with feedback becomes indistinguishable from that of the system without feedback (i.e. Δlog(*Y*) = Δlog(*Y*_*NF*_)). This procedure can be repeated for any parameter set of interest. Specifically, we can compute CoRa for a range of values of the parameter *θ* ∈ Θ while adjusting the constant input of the no-feedback system (as explained above) accordingly to ensure the mathematically controlled comparison in the sense we describe above. We therefore uniformly use the notation CoRa_*θ*∈Θ_(*ρ*) to designate the CoRa function computed for a perturbation in *ρ* when the system is positioned at some changing value of parameter *θ* ∈ Θ. CoRa_*θ*∈Θ_(*ρ*) is therefore a representation of the capacity of feedback to mediate adaptation of the system’s output to perturbations to the parameter *ρ* for every value of *θ* considered. In this work, *θ* is limited to a change in an individual parameter.

## Using CoRa to characterize negative feedback in a system architecture capable of perfect adaptation

We first tested CoRa on a well-established negative feedback control structure, the antithetic feedback motif, which can exhibit perfect adaptation to step disturbance inputs when connected to an arbitrarily complex biochemical network [2] (Fig. 2A). The antithetic motif is composed of two molecular species that annihilate each other through their mutual binding. One of the antithetic molecular species controls the input of a biochemical network and the other is produced by the output of the same network. If the antithetic molecules are only lost through the mutual annihilation event without individual degradation or dilution, this strategy is expected to generate a system with perfect adaptation to a step perturbation [2]. Using CoRa to study this feedback motif, we recapitulate this result, showing that perfect adaptation is possible (Fig. 2). Interestingly, our analysis also reveals that relaxing the assumption of zero dilution and adding molecular details such as explicit accounting of the transitory molecule resulting from binding of the two antithetic molecules (complex *C* in Fig. 2A) is sufficient to compromise perfect adaptation, often in non-trivial ways (Fig. 2B-C). For example, 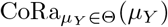 (*μ*_*Y*_ is the synthesis rate of the output molecule *Y*) deviates from perfect adaptation value of 0 if dilution of antithetic molecules is assumed to occur individually at a small rate *γ* = 10^−4^ *min*^−1^ (Fig. 2B,D). This deviation from perfect adaptation occurs at low and high values of *μ*_*Y*_ (Fig. 2B; *Supplementary Information*). In a further elaboration of the circuit, when we consider the complex *C* as a functional molecule that can influence the synthesis of the output molecule *Y* until its removal from the system [11] (Fig. 2A), the feedback undergoes a dramatic failure in its ability to produce perfect adaptation after a specific threshold value of *μ*_*Y*_. This is evidenced by 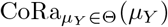 shifting abruptly from almost zero to one (Fig. 2B,D). This is also the case for 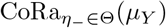 (*η*_−_ is degradation rate of complex *C*; Fig. 2C). These conclusions are not unique to the particular controlled subsystem considered (Fig. S2), or the perturbation used (Fig. S3). Exploration of this phenomenology identified by CoRa reveals that this qualitative change in the feedback control results from saturation in the concentration of the complex *C* (*Supplementary Information*; Fig. S4), an insight that would have been difficult without the computational observation of this behavior.

**Figure 2.**
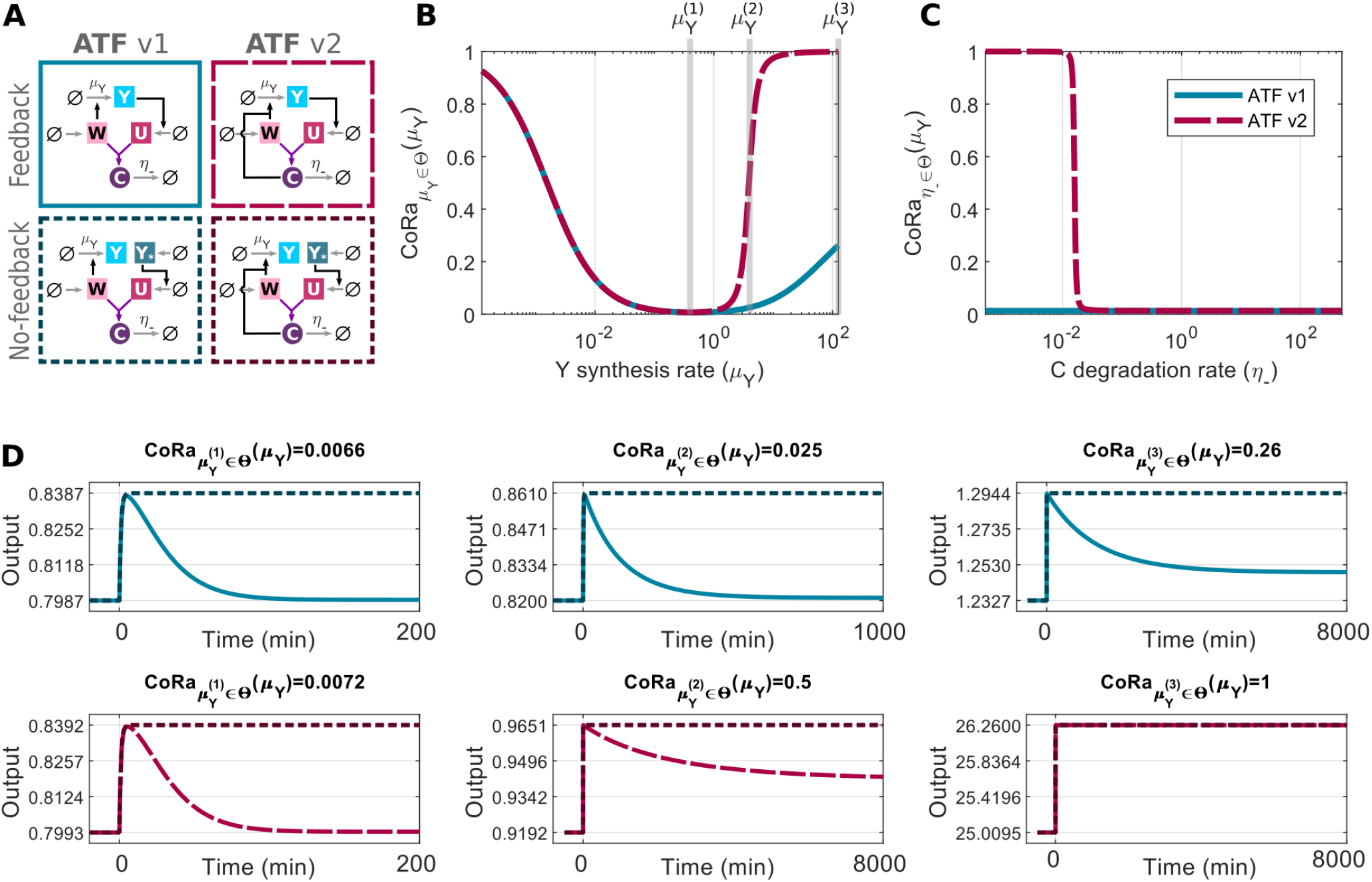
Characterizing the Antithetic Feedback Motif (ATF) using CoRa. **(A)** The ATF motif is composed by two molecules (*W, U*) that bind and inactivate each other, forming a transitory complex *C* which is then degraded with rate *η*_−_; one antithetic molecule *W* induces *Y* synthesis (the controlled output, with rate *μ*_*Y*_), and *Y* feeds back by inducing the synthesis of the other antithetic molecule, *U*. We consider two variations of the feedback structure. The first (ATF v1; blue continuous lines) is akin to the original ATF motif with the difference that the binding of *U* and *W* generates a complex *C* that is explicitly modeled before it disappears through degradation at a rate *η*_−_. In the second feedback structure (ATF v2; pink long-dash lines), the complex *C* retains biological activity in influencing the production of *Y* until it is degraded. This structure is inspired by the feedback implementation documented in Ng *et al*. [11]. For each case, the associated locally analogous system without feedback (bottom box with dotted line) is shown. **(B)** CoRa computed following perturbations to *μ*_*Y*_, the synthesis rate of *Y*, as this parameter itself is varied. **(C)** CoRa computed following a perturbation to *μ*_*Y*_, the synthesis rate of *Y*, as the degradation rate of the complex *C* (*η*_−_) is varied. **(D)** The output response of the ATF system (output is *Y*, blue continuous lines, v1, and pink long-dash lines, v2) and associated locally analogous systems without feedback (dark dash lines) as a function of time after a small perturbation occurs on *μ*_*Y*_ (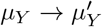 at time zero). Plots are shown for three different *Y* synthesis rate values 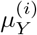 (highlighted in panel B with gray vertical lines). The resulting 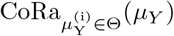 value is shown as the plot title for each case. See *Supplementary Information* for equations and Table S1 for parameter values.

## Using CoRa to compare different feedback control mechanisms

Any feedback control system can be analyzed using CoRa, providing a unifying framework under which different feedback mechanisms can be rigorously compared. In fact, we were able to rapidly analyze a large number of distinct feedback control motifs proposed in the literature [2, 4, 6, 7, 11, 14, 16] (Fig. 3; *Supplementary Information*). For comparison on equal footing, we considered each of these different negative feedback structures controlling the same simple biochemical subsystem. These investigations using CoRa generated a rich data-set to explore the properties of different molecular implementations associated with the same phenomenological macroscopic function –negative auto-regulation. For example, it was clear that specific molecular details of feedback generate distinct adaptation properties to the same perturbation. As only one example, 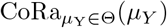 computed for all feedback strategies employing repression of synthesis modelled using a standard Michaelis-Menten repression function (Fig. 3G-I) displayed a limit of CoRa_*θ*∈Θ_(*μ*_*Y*_) ≥ 0.5. This behavior relates to the inevitable saturation of the repression function (see *Supplementary Information* for an example of an analytical treatment of this limit). A notable exception to this limit occurred for the “brink motif” feedback strategy, a motif that combines antithetic molecular sequestration with an activation-deactivation enzymatic cycle to produce a tuneable ultra-sensitive response [14] (Fig. 3K). These patterns that were computationally pinpointed by the CoRa analysis prompted the hypothesis that adding ultra-sensitivity to motifs with Michaelis-Menten synthesis repression might alleviate the limit of their adaptive behaviors. Using CoRa, we tested this hypothesis by adding a Hill coefficient larger than 1 to the Michaelis-Menten function in different strategies. By increasing the system ultrasensitivity with the Hill coefficient, the lower bound of the CoRa curve decreased in all cases, indicating improved adaptation capabilities of the control loops (Fig. S5A-C). Furthermore, increasing the ultrasensitivity of the brink motif itself by increasing the deactivation rate in its enzymatic cycle [14] improved its ability to adapt (Fig. S5D). These results strongly suggest that feedback strategies based on Michaeliean repression of synthesis are severely limited in their capacity for homeostasis, but can be improved using ultra-sensitive components. In this case, CoRa was used as a computational hypothesis generator about this general principle, which was then confirmed through further computational and analytical investigations. Finally, identification of strategies for improving feedback performance can be automated by embedding CoRa into an optimization framework in which the parameters of the feedback are iteratively changed to generate a desirable CoRa curve (Fig. S6). This optimization procedure can help uncover the parameter constraints that are needed for adaptation given the specific biochemical feedback structure in a system. It can also help in the design of *de novo* synthetic feedback structures in cells.

**Figure 3.**
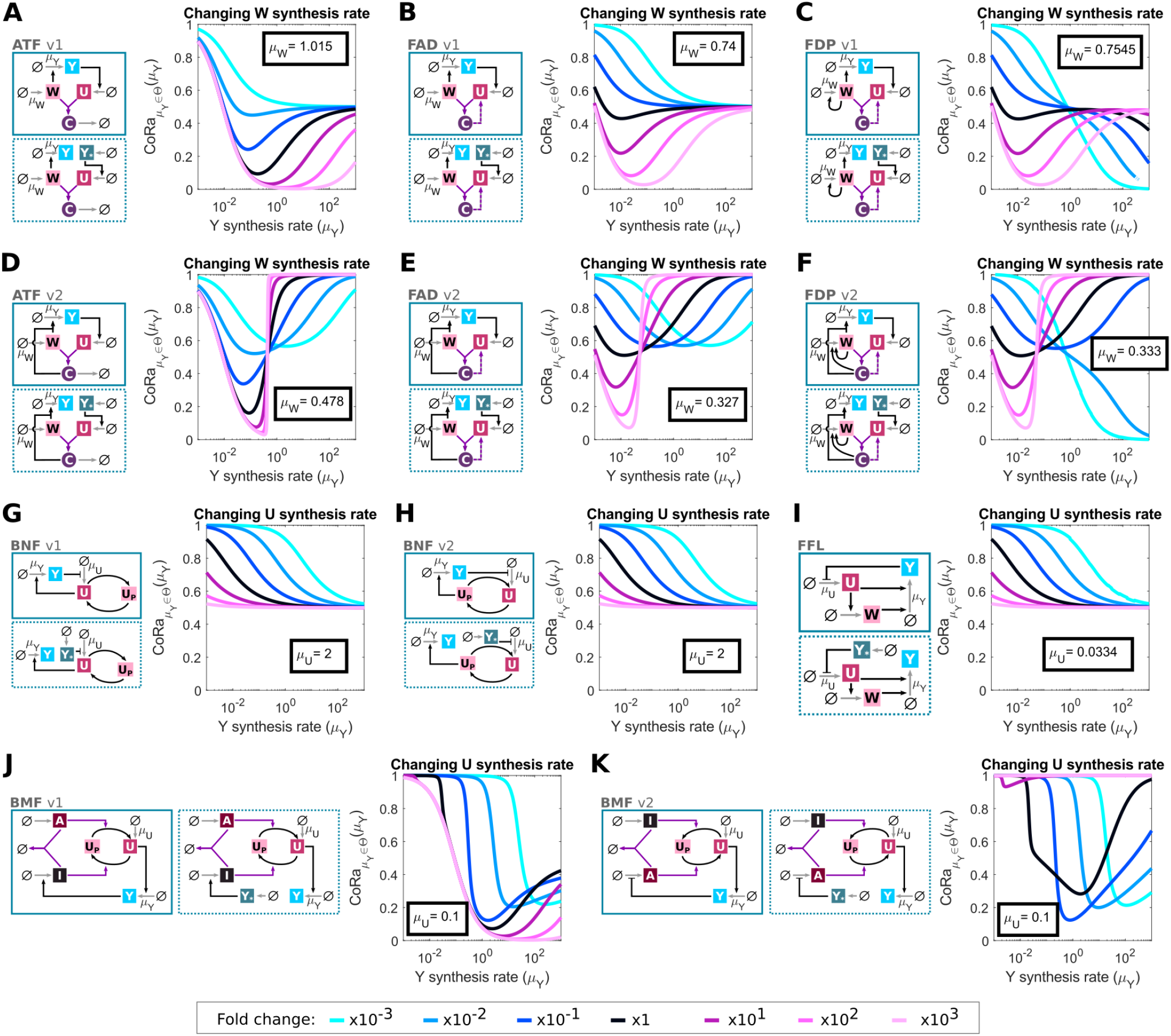
CoRa provides a unifying framework to compare different feedback control architectures. (A-K) Different feedback motifs, with different levels of complexity, can be directly compared using the CoRa function. In each case, the diagrams of the feedback system (in box with continuous line) and its associated locally analogous system without feedback (in box with dotted line) are shown. 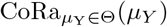 is computed for 7 different values of a given parameter that is also varied in addition to *μ*_*Y*_. The identity and nominal value of the varied parameter (either *μ*_*W*_ the *W* synthetic rate, or *μ*_*U*_ the *U* synthesis rate) is indicated on every plot, and how it is varied is shown at the bottom of the figure with appropriate color-coding information. Different mechanisms show different signature behaviors; particularly, for ATF-like architectures, a complex *C* of the two antithetic molecules that retains activity results in an abrupt loss of control (D-F) compared to the analogous motifs with an inactive complex (A-C). The Michaelis-Menten function describing feedback through repression of synthesis imposes a limit on the efficiency of feedback (G-I), except in the presence of ultrasensitivity (K). ATF: Antithetic Feedback, FAD: Feedback by Active Degradation, FDP: Feedback by Active Degradation + Positive Feedback, BNF: Buffering + Negative Feedback, FFL: Negative Feedback + Feed-forward loop, and BMF: Brinf Motif. In all cases, the controlled subsystem parameters are identical, and the output steady state *Y* ≈ 10 *nM* for the “× 1” case (black line) and *μ*_*Y*_ = 1 *min*^−1^. See *Supplementary Information* for equations and Table S1 for parameter values.

## Discussion

A framework for the systematic evaluation and comparison of biochemical feedback control systems is essential for understanding the general principles of biological homeostasis. While many methods exist for the evaluation of technological feedback systems, understanding the principles of biological adaptation mediated through feedback poses its unique challenges, including distinct mathematical properties of the biological substrate. Importantly, the nature of biological organization with extensive coupling of parameters and processes makes the extraction of engineering-centric quantities needed for traditional analyses of feedback quantities, such as setpoints and regulation errors, challenging. Debate about whether these quantities are defined for biological systems has a long history and no concrete resolution [13]. One advantage of CoRa is that it does not make any assumptions about the existence of such quantities, replacing this debate with a comparison to a system that would have evolved identically but without the feedback structure. Another advantage of CoRa is that it is agnostic to the complexity of the system. While we have only used a simple system to illustrate the properties of CoRa, extending analysis to more complex systems was straightforward (Fig. S2). We have also used CoRa to assess feedback-mediated adaptation for only one perturbation as a function of one model parameter. However, it should be easy to see that a multi-dimensional CoRa for simultaneous perturbations or many concurrent parameter changes is easily computable. We expect, however, that new methods would be needed to analyze the resulting multi-dimensional data into coherent principles.

Finally, the concept of CoRa should be easily extendable to assessing the quantitative contribution of feedback to important properties other than the steady-state response to perturbation. These might, for example, include the role of feedback in the dynamic response of a system or its response to stochastic fluctuations. As such, CoRa represents a flexible framework that is poised to catalyze fast progress in our understanding of the many roles that feedback control plays in biological organization.

## SUPPLEMENTARY MATERIALS

**Supplementary Figures:** Fig S1-S6

**Supplementary Tables:** Table S1-S2

**Supplementary Text:** Section S1-S5

**Supplementary Code:** https://github.com/mgschiavon/CoRa/releases/tag/v1.0

## Data availability

The authors declare that all the data supporting the findings of this study are available within the article and its Supplementary Information files.

## Code availability

The computer code used to generate and analyze the data in this study is available at https://github.com/mgschiavon/CoRa/releases/tag/v1.0.

## Acknowledgments

The authors would like to thank the members of the El-Samad lab for useful comments and discussion on the manuscript. This work was supported by the Chan-Zuckerberg Biohub and the Defense Advanced Research Projects Agency, Contract No. HR0011-16-2-0045 to H.E.-S, and by ANID—Millennium Science Initiative Program—Millennium Institute for Integrative Biology (iBio ICN17 022) to M.G.-S. The content and information does not necessarily reflect the position or the policy of the government, and no official endorsement should be inferred. H.E.-S. is a Chan-Zuckerberg investigator.

## SUPPLEMENTARY INFORMATION

**Figure S1.**
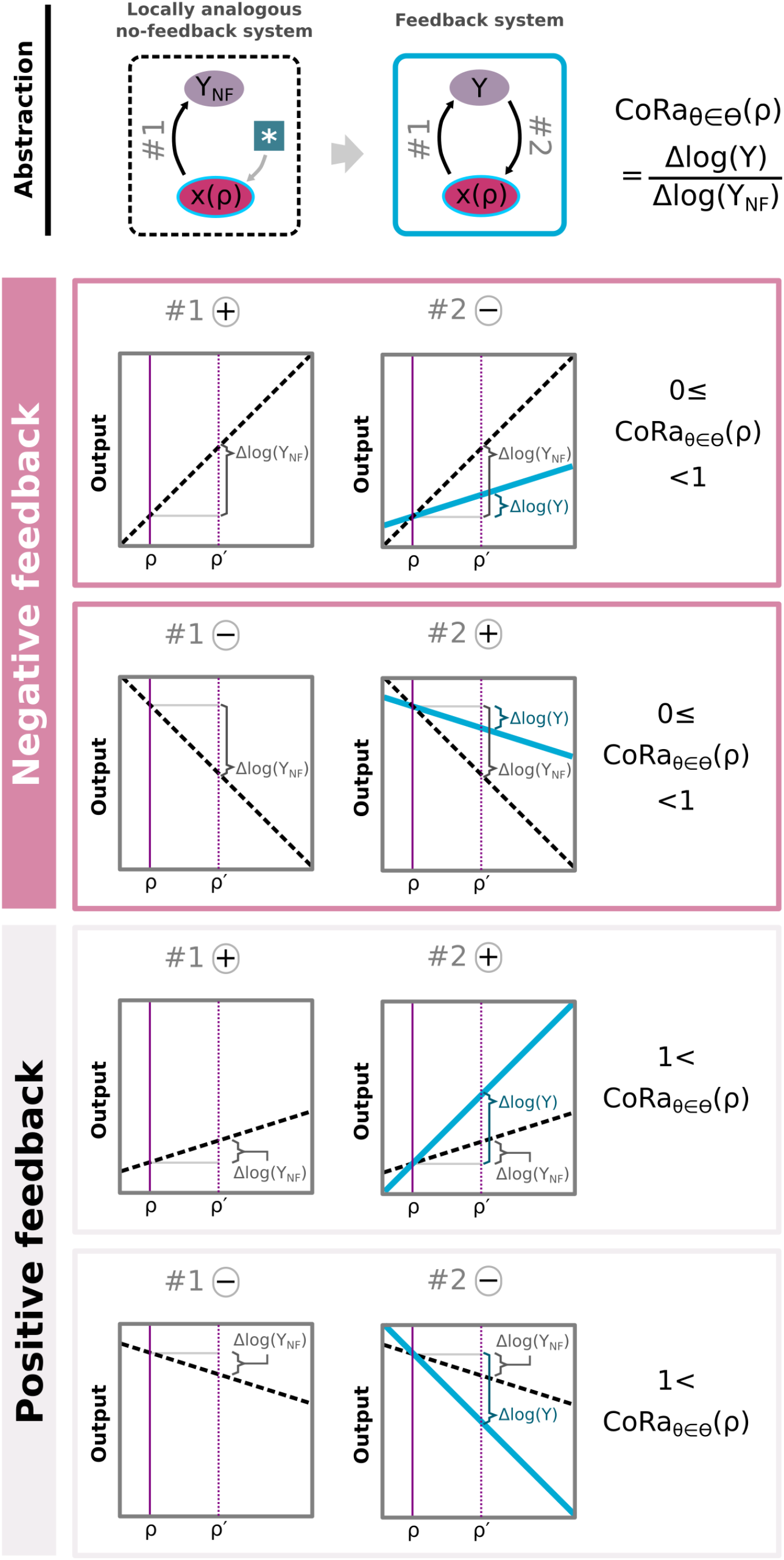
Feedback logic and CoRa values. We abstract the control system to a two-node network where one node represents the output to be controlled (*Y*), and the other the rest of the system including the dependency on the parameter *ρ* to be perturbed (*x*(*ρ*)). The locally analogous system can be represented as an equivalent network, with a third node (*) that represent the new input into the *x*(*ρ*) node. The other link from *x*(*ρ*) to the output (*Y*_*NF*_ ; link #1) remains the same between the two networks. (Left column) The sign of link #1 can be determined by comparing the output before (*Y*_*NF*_|_Θ_ = *Y*|_Θ_) and after (*Y*_*NF*_ | _Θ,*ρ*→*ρ*′_) the perturbation. For a positive perturbation, link #1 is positive (#1 (+)) if and only if *Y*_*NF*_|_Θ,*ρ*→*ρ*′_ > *Y*, or negative (#1 (−)) if and only if *Y*_*NF*_ | _Θ,*ρ*→*ρ*′_ < *Y*. (Middle column) The sign of the feedback link from the output to the *x*(*ρ*) node (link #2) can be determined by comparing the output after the perturbation in the feedback system (*Y*|_Θ,*ρ*→*ρ*′_) and in the locally analogous system (*Y*_*NF*_| _Θ,*ρ*→*ρ* ′_). It is positive (#2 (+)) if and only if *Y*|_Θ,*ρ*→*ρ*′_ > *Y*_*NF*_|_Θ,*ρ*→*ρ* ′_, or negative (#2 (−)) if and only if *Y*|_Θ,*ρ*→*ρ*′_ < *Y*_*NF*_ | _Θ,*ρ*→*ρ* ′_. (Right column) Given the formula for CoRa, we can see that CoRa_*θ*∈Θ_(*ρ*) is bound between 0 and 1 whenever we have a negative feedback, and bigger than 1 in the case of a positive feedback.

**Figure S2.**
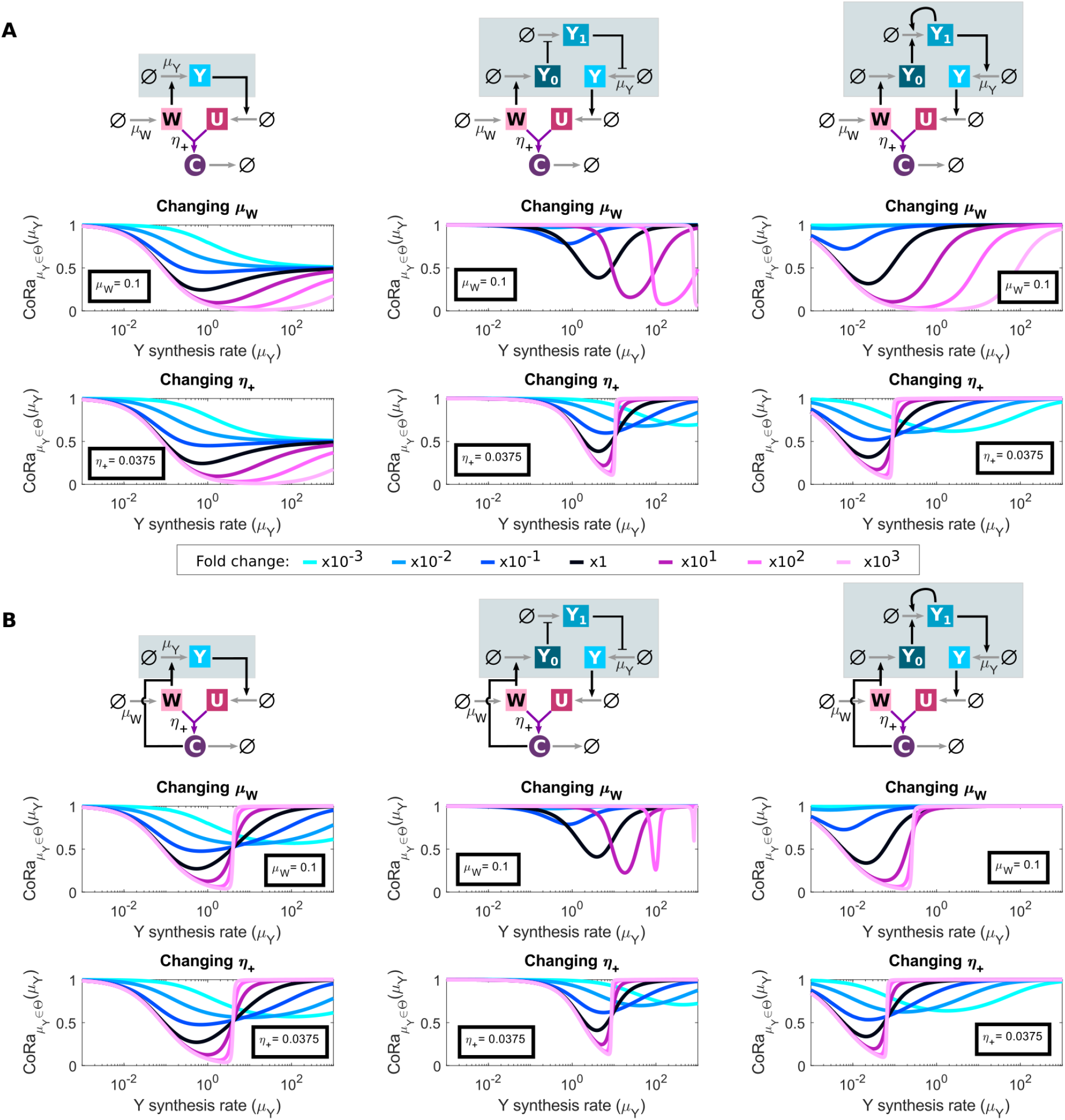
Antithetic feedback control performance depends on controlled subsystem. Three different subsystems (Eqs. S38-S40) of increasing complexity (controlled subsystem highlighted in gray) controlled by the antithetic feedback control (ATF) can be compared using the CoRa function. **(A)** CoRa plots for modified ATF with inactive complex *C* (v1). First row shows a schematic controlled subsystem. 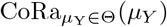 is computed for 7 different values of a given parameter that is also varied in addition to *μ*_*Y*_. The identity and nominal value of the varied parameter (either *μ*_*W*_ the *W* synthetic rate, or *η*_+_ the *U* : *W* binding rate) is indicated on every plot, and how it is varied is shown in between the two panels of the figure with appropriate color-coding information. **(B)** Same as **(A)** but modified ATF with active complex *C* (v2). See Section S4 for equations and Table S2 for parameter values.

**Figure S3.**
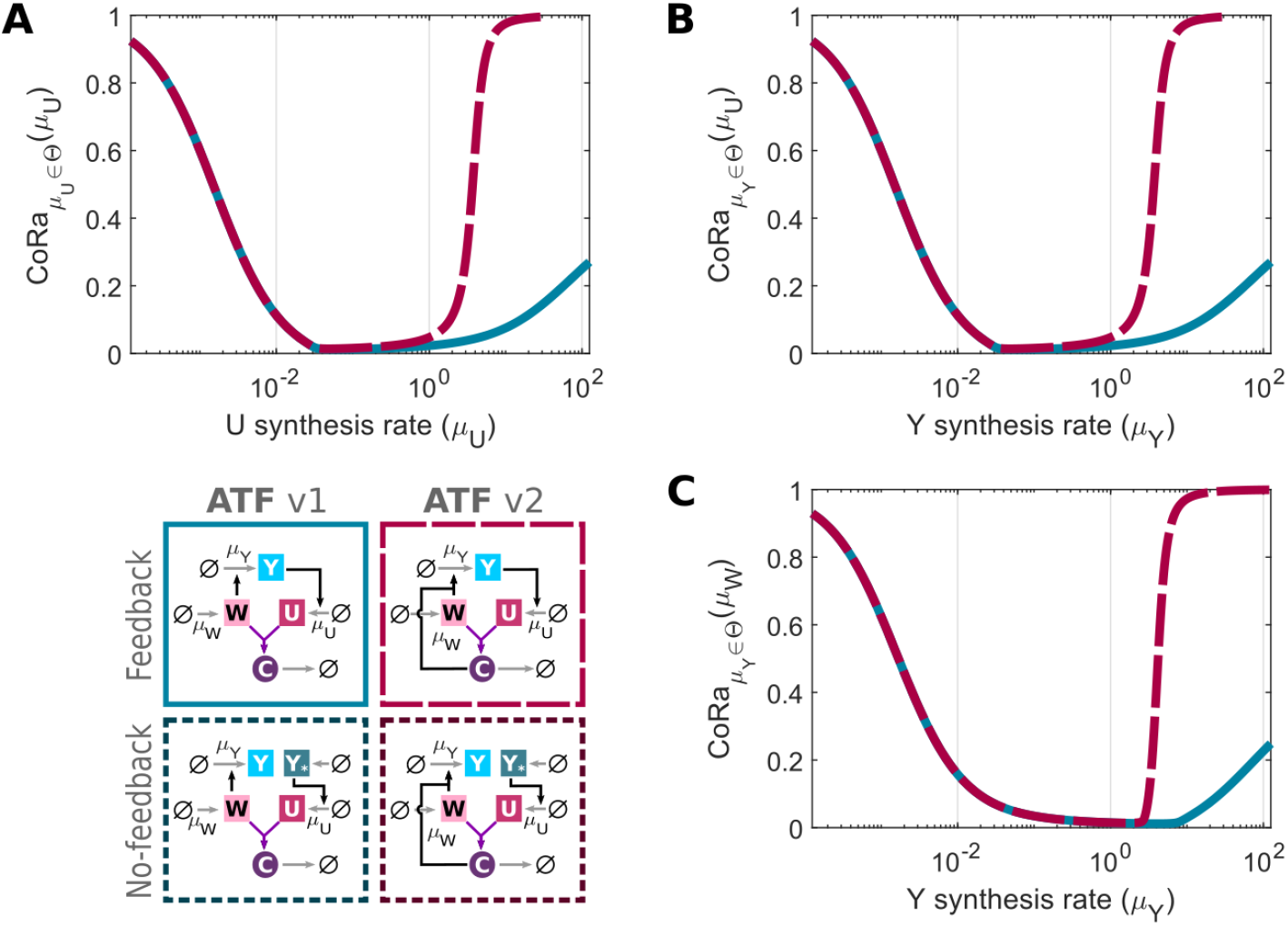
CoRa can be computed for perturbation of any parameter as a function of another parameters. Plots are shown for the two versions of the modified antithetic feedback (ATF) control. **(A)** CoRa plot as a function of *U* synthesis rate (*μ*_*U*_) as *μ*_*U*_ itself is perturbed. **(B)** CoRa plot as a function of *Y* synthesis rate (*μ*_*Y*_) as *μ*_*U*_ is perturbed. **(C)** CoRa plot as a function of *Y* synthesis rate (*μ*_*Y*_) as *W* synthesis rate (*μ*_*W*_) is perturbed. ATF v1, blue continuous lines; ATF v2, pink long-dash lines. See Section S4 for equations and Table S2 for parameter values.

**Figure S4.**
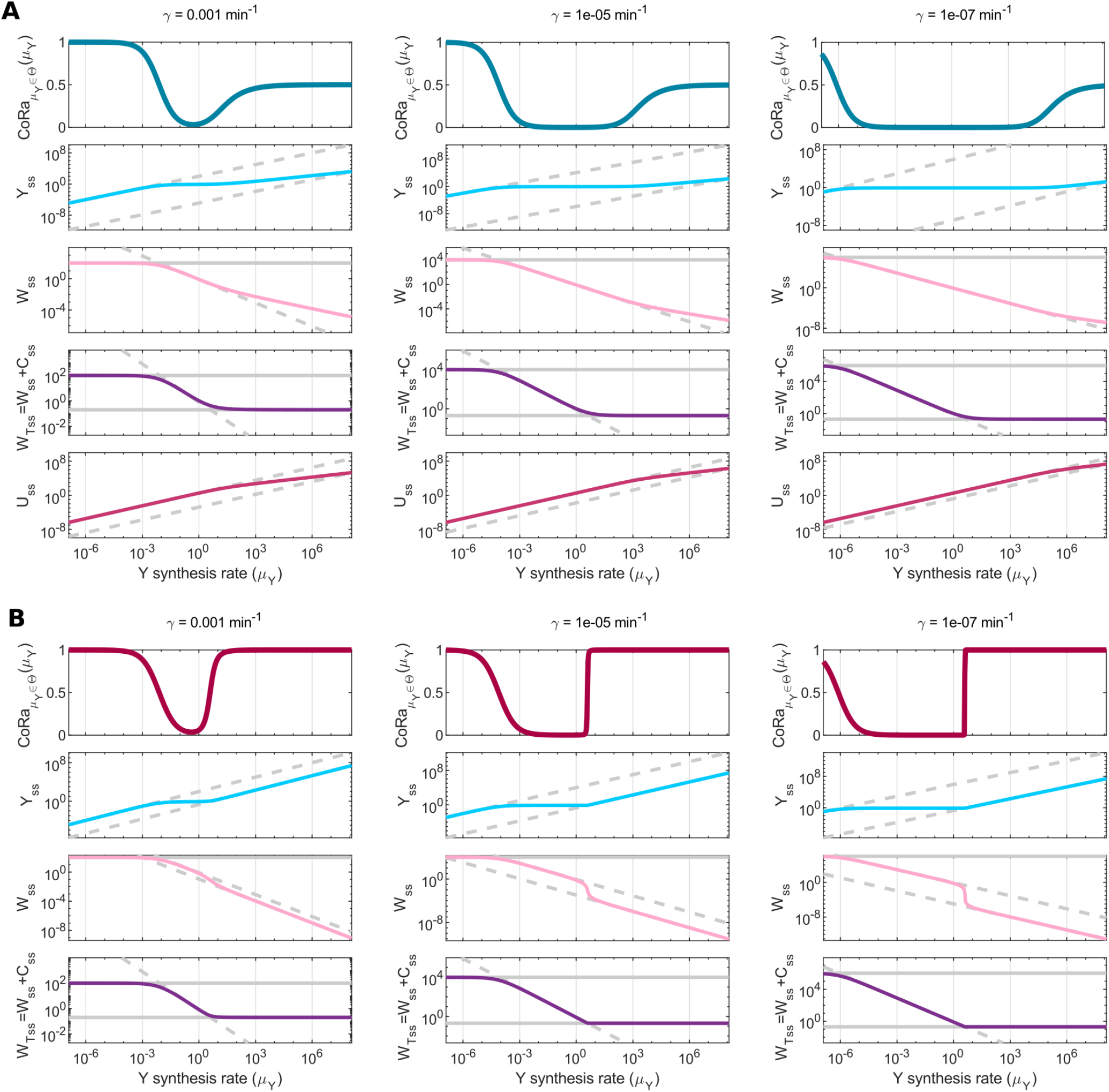
Effect of dilution on the modified antithetic feedback (ATF) control and system saturation. Effect of dilution (*γ*; see column titles) on the ATF control performance following perturbations to *μ*_*Y*_, the synthesis rate of *Y*, as this parameter itself is varied, with either inactive (v1; **A**) or active (v2; **B**) complex *C*. **(A)** For ATF v1, as *μ*_*Y*_ decreases, 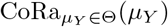 increases. When *W* steady state concentration (*W*_*ss*_) saturates approaching its limit value 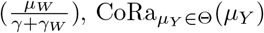 approaches 1. On the other extreme, as *μ*_*Y*_ increases, 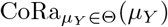 increases. When total *W* at steady-state (*W*_*T,ss*_ = *W*_*ss*_ + *C*_*ss*_) concentration saturates 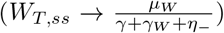, *U* steady-state concentration (*U*_*ss*_) cannot increase proportionally to *μ*_*Y*_ to allow free *W*_*ss*_ to decrease in the same proportion (given that in steady state, 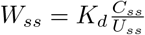; see Section S2.1.1). **(B)** For ATF v2, 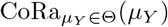 increases for both low and high *μ*_*Y*_ values as total *W* steady state concentration (*W*_*T,ss*_ = *W*_*ss*_ + *C*_*ss*_) saturates, reaching its higher 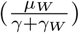 and lower 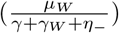 limit values, respectively (see Section S2.1.2). In all plots, limits are shown as horizontal gray lines, and gray dashed lines increasing or decreasing proportionally to *μ*_*Y*_ are shown as reference. See Section S4 for equations and Table S2 for parameter values.

**Figure S5.**
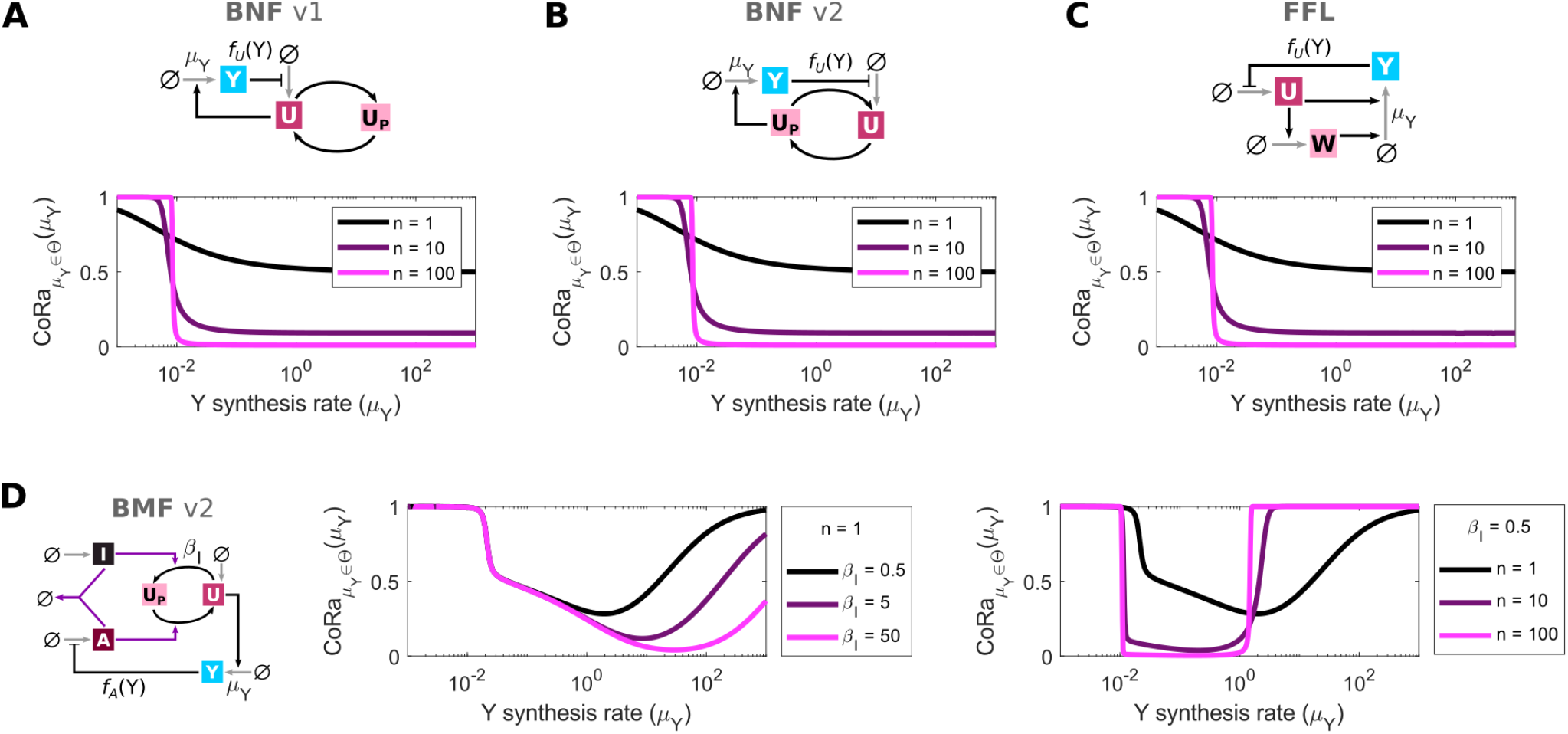
Negative auto-regulation affecting synthesis represented by Michaelis-Menten function limits control performance in multiple motifs, but is alleviated by ultrasensitivity. In this figure, the negative auto-regulation function is modeled as a negative Hill function, 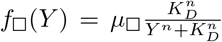, where *μ*□ is the maximum synthesis rate, *K*_*D*_ is the *EC*_50_, and *n* is the Hill coefficient. Four of the explored motifs in Fig. 3 include negative synthesis regulation (*f*_□_(*Y*)): **(A-B)** Buffering + Negative Feedback (BNF v1 & v2; Fig. 3G-H), **(C)** Feedback + Feedforward Loop (FFL; Fig. 3I), and **(D)** Brink Motif Feedback with repression of activator (BMF v2; Fig. 3K). For each motif, plots show CoRa function for perturbations to the *Y* synthesis rate (*μ*_*Y*_) as the Hill coefficient *n* increases. (In all cases, the black line corresponds to the black line in Fig. 3). For **(D)** BMF v2, we also show how the CoRa function changes while increasing the inactivation rate *β*_*I*_ ([*nM* ^−1^ *min*^−1^]) from *U* to *U*_*P*_, which is dependent on *I*. We corroborate that, as shown by Samaniego & Franco [14], the BMF motif displays high ultrasensitivity, and the ultrasensitivity increases as *β*_*I*_ increases. In all cases, higher ultrasensitivity (either by increasing the Hill coefficient *n* or *β*_*I*_ for BMFv2) results in improved control performance for some range of *μ*_*Y*_ values (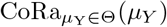 approaching zero). See Section S4 for equations and Table S2 for parameter values.

**Figure S6.**
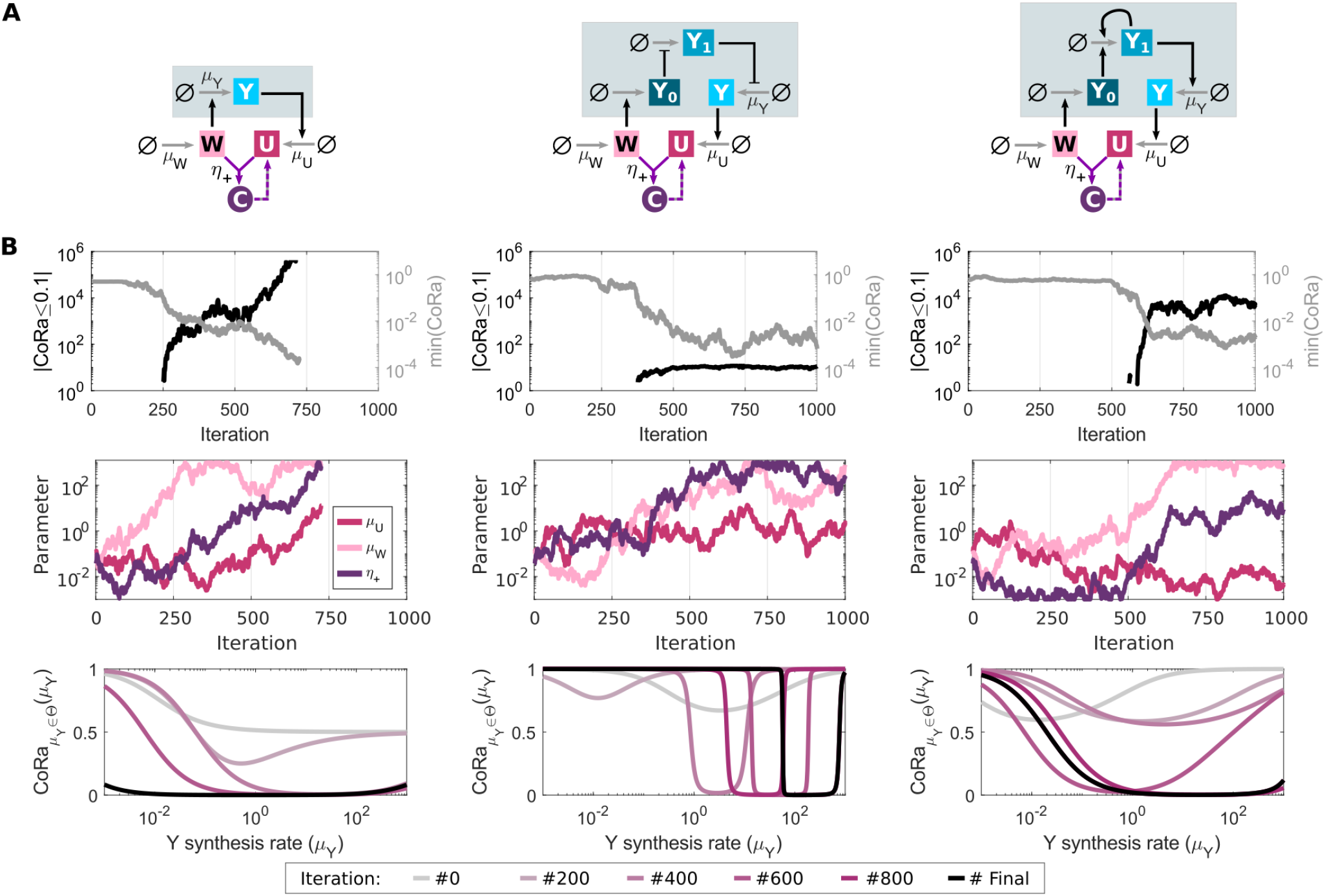
Optimizing a controller for different subsystems. **(A)** Diagrams for three different subsystems (Eqs. S38-S40; gray boxes) that are controlled using the feedback by active degradation motif (FAD v1). **(B)** The feedback control parameters (*U* synthesis rate dependent on *Y, μ*_*U*_ ; *W* constitutive synthesis rate, *μ*_*W*_ ; and *U, W* binding rate, *η*_+_) can be optimized for each subsystem to drive CoRa below a given threshold 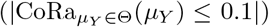 for a large dynamic range in *μ*_*Y*_, the synthesis rate of *Y*. The optimization stops after 1000 iterations or whenever 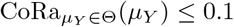 for the whole range of *μ*_*Y*_ values considered; see *Section* S5.1 for algorithm details. Optimization traces 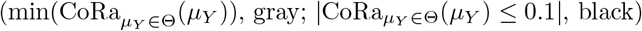, as well as the associated parameter values ({ *μ*_*U*_, *μ*_*W*_, *η*_+_}), are shown for each system; the 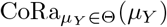 curves for some iterations are also shown. See Section S4 for equations and Table S2 for parameter values.

**Table S1.**
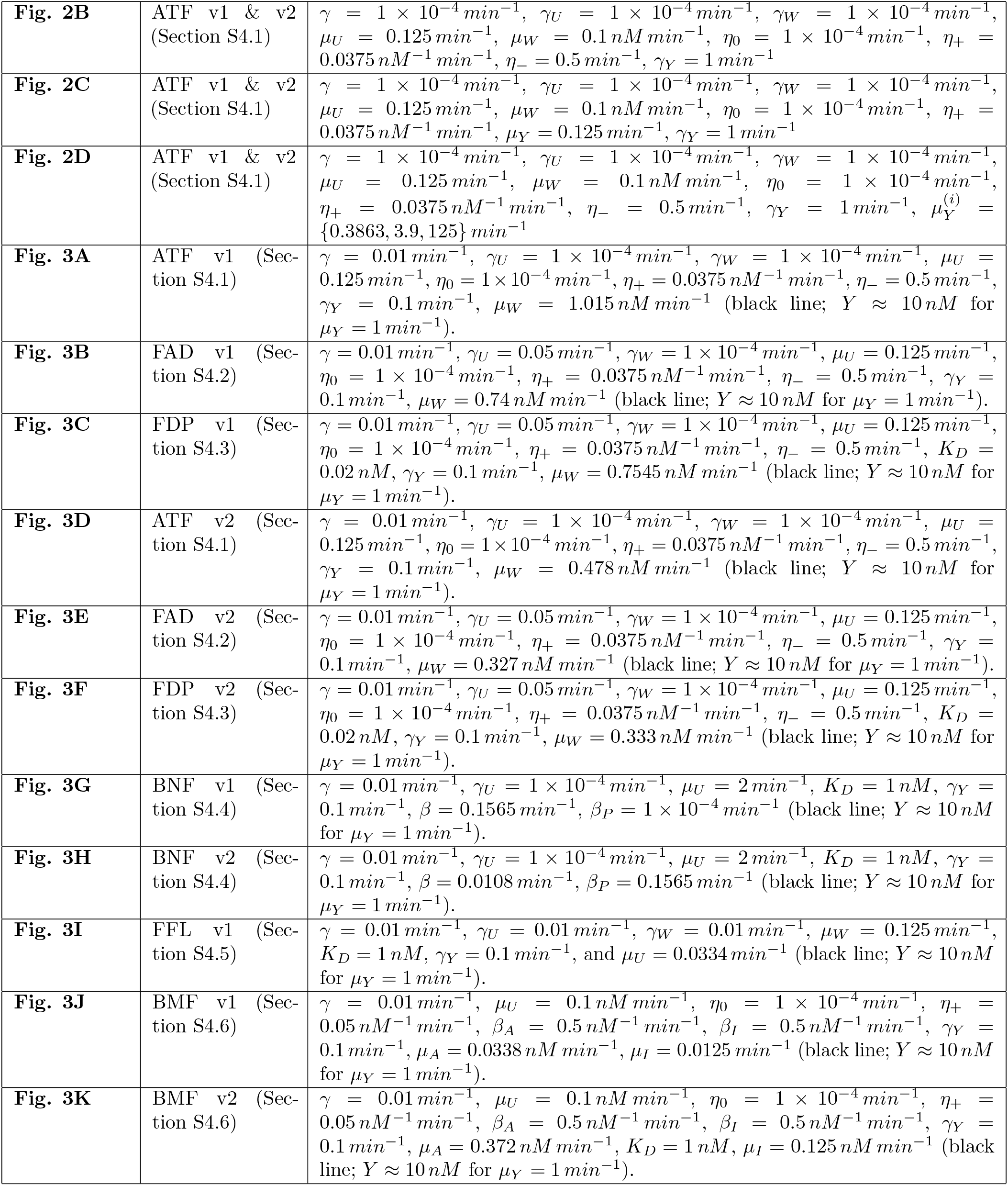
Used parameter values in main figures.

**Table S2.**
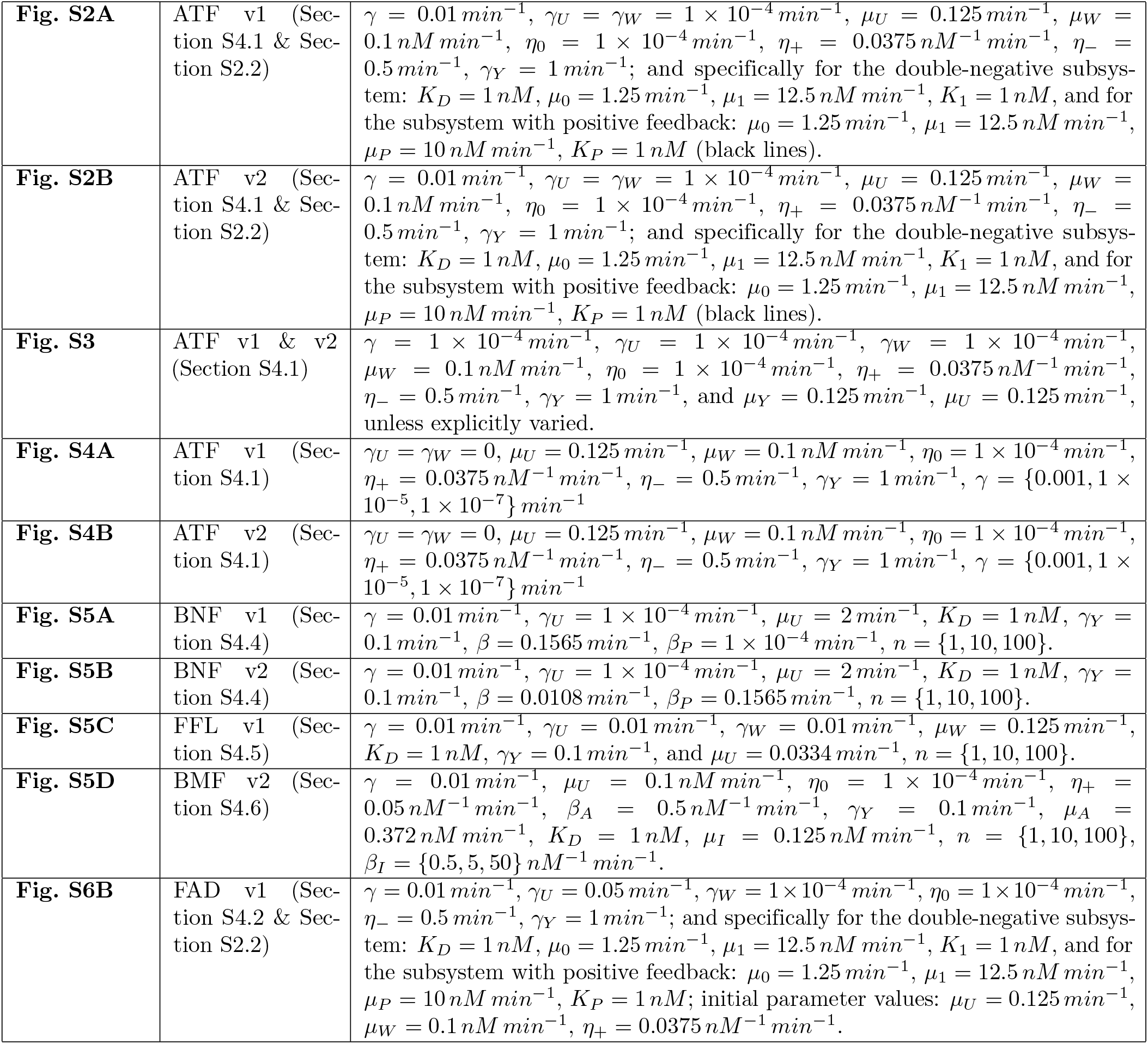
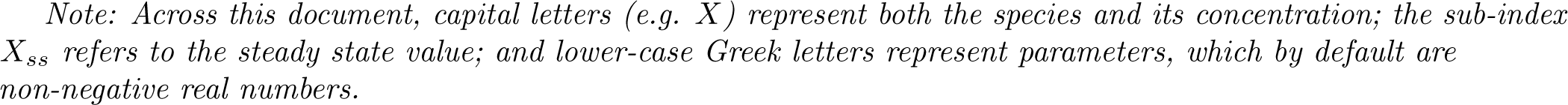
Used parameter values in supplementary figures.

|  |  |  |
| --- | --- | --- |
| **Fig. S2A** | ATF v1 (Section S4.1 & Section S2.2) | $\gamma = 0.01 \text{ min}^{-1}$, $\gamma_U = \gamma_W = 1 \times 10^{-4} \text{ min}^{-1}$, $\mu_U = 0.125 \text{ min}^{-1}$, $\mu_W = 0.1 \text{ nM min}^{-1}$, $\eta_0 = 1 \times 10^{-4} \text{ min}^{-1}$, $\eta_+ = 0.0375 \text{ nM}^{-1} \text{ min}^{-1}$, $\eta_- = 0.5 \text{ min}^{-1}$, $\gamma_Y = 1 \text{ min}^{-1}$; and specifically for the double-negative subsystem: $K_D = 1 \text{ nM}$, $\mu_0 = 1.25 \text{ min}^{-1}$, $\mu_1 = 12.5 \text{ nM min}^{-1}$, $K_1 = 1 \text{ nM}$, and for the subsystem with positive feedback: $\mu_0 = 1.25 \text{ min}^{-1}$, $\mu_1 = 12.5 \text{ nM min}^{-1}$, $\mu_P = 10 \text{ nM min}^{-1}$, $K_P = 1 \text{ nM}$ (black lines). |
| **Fig. S2B** | ATF v2 (Section S4.1 & Section S2.2) | $\gamma = 0.01 \text{ min}^{-1}$, $\gamma_U = \gamma_W = 1 \times 10^{-4} \text{ min}^{-1}$, $\mu_U = 0.125 \text{ min}^{-1}$, $\mu_W = 0.1 \text{ nM min}^{-1}$, $\eta_0 = 1 \times 10^{-4} \text{ min}^{-1}$, $\eta_+ = 0.0375 \text{ nM}^{-1} \text{ min}^{-1}$, $\eta_- = 0.5 \text{ min}^{-1}$, $\gamma_Y = 1 \text{ min}^{-1}$; and specifically for the double-negative subsystem: $K_D = 1 \text{ nM}$, $\mu_0 = 1.25 \text{ min}^{-1}$, $\mu_1 = 12.5 \text{ nM min}^{-1}$, $K_1 = 1 \text{ nM}$, and for the subsystem with positive feedback: $\mu_0 = 1.25 \text{ min}^{-1}$, $\mu_1 = 12.5 \text{ nM min}^{-1}$, $\mu_P = 10 \text{ nM min}^{-1}$, $K_P = 1 \text{ nM}$ (black lines). |
| **Fig. S3** | ATF v1 & v2 (Section S4.1) | $\gamma = 1 \times 10^{-4} \text{ min}^{-1}$, $\gamma_U = 1 \times 10^{-4} \text{ min}^{-1}$, $\gamma_W = 1 \times 10^{-4} \text{ min}^{-1}$, $\mu_W = 0.1 \text{ nM min}^{-1}$, $\eta_0 = 1 \times 10^{-4} \text{ min}^{-1}$, $\eta_+ = 0.0375 \text{ nM}^{-1} \text{ min}^{-1}$, $\eta_- = 0.5 \text{ min}^{-1}$, $\gamma_Y = 1 \text{ min}^{-1}$, and $\mu_Y = 0.125 \text{ min}^{-1}$, $\mu_U = 0.125 \text{ min}^{-1}$, unless explicitly varied. |
| **Fig. S4A** | ATF v1 (Section S4.1) | $\gamma_U = \gamma_W = 0$, $\mu_U = 0.125 \text{ min}^{-1}$, $\mu_W = 0.1 \text{ nM min}^{-1}$, $\eta_0 = 1 \times 10^{-4} \text{ min}^{-1}$, $\eta_+ = 0.0375 \text{ nM}^{-1} \text{ min}^{-1}$, $\eta_- = 0.5 \text{ min}^{-1}$, $\gamma_Y = 1 \text{ min}^{-1}$, $\gamma = \{0.001, 1 \times 10^{-5}, 1 \times 10^{-7}\} \text{ min}^{-1}$ |
| **Fig. S4B** | ATF v2 (Section S4.1) | $\gamma_U = \gamma_W = 0$, $\mu_U = 0.125 \text{ min}^{-1}$, $\mu_W = 0.1 \text{ nM min}^{-1}$, $\eta_0 = 1 \times 10^{-4} \text{ min}^{-1}$, $\eta_+ = 0.0375 \text{ nM}^{-1} \text{ min}^{-1}$, $\eta_- = 0.5 \text{ min}^{-1}$, $\gamma_Y = 1 \text{ min}^{-1}$, $\gamma = \{0.001, 1 \times 10^{-5}, 1 \times 10^{-7}\} \text{ min}^{-1}$ |
| **Fig. S5A** | BNF v1 (Section S4.4) | $\gamma = 0.01 \text{ min}^{-1}$, $\gamma_U = 1 \times 10^{-4} \text{ min}^{-1}$, $\mu_U = 2 \text{ min}^{-1}$, $K_D = 1 \text{ nM}$, $\gamma_Y = 0.1 \text{ min}^{-1}$, $\beta = 0.1565 \text{ min}^{-1}$, $\beta_P = 1 \times 10^{-4} \text{ min}^{-1}$, $n = \{1, 10, 100\}$. |
| **Fig. S5B** | BNF v2 (Section S4.4) | $\gamma = 0.01 \text{ min}^{-1}$, $\gamma_U = 1 \times 10^{-4} \text{ min}^{-1}$, $\mu_U = 2 \text{ min}^{-1}$, $K_D = 1 \text{ nM}$, $\gamma_Y = 0.1 \text{ min}^{-1}$, $\beta = 0.0108 \text{ min}^{-1}$, $\beta_P = 0.1565 \text{ min}^{-1}$, $n = \{1, 10, 100\}$. |
| **Fig. S5C** | FFL v1 (Section S4.5) | $\gamma = 0.01 \text{ min}^{-1}$, $\gamma_U = 0.01 \text{ min}^{-1}$, $\gamma_W = 0.01 \text{ min}^{-1}$, $\mu_W = 0.125 \text{ min}^{-1}$, $K_D = 1 \text{ nM}$, $\gamma_Y = 0.1 \text{ min}^{-1}$, and $\mu_U = 0.0334 \text{ min}^{-1}$, $n = \{1, 10, 100\}$. |
| **Fig. S5D** | BMF v2 (Section S4.6) | $\gamma = 0.01 \text{ min}^{-1}$, $\mu_U = 0.1 \text{ nM min}^{-1}$, $\eta_0 = 1 \times 10^{-4} \text{ min}^{-1}$, $\eta_+ = 0.05 \text{ nM}^{-1} \text{ min}^{-1}$, $\beta_A = 0.5 \text{ nM}^{-1} \text{ min}^{-1}$, $\gamma_Y = 0.1 \text{ min}^{-1}$, $\mu_A = 0.372 \text{ nM min}^{-1}$, $K_D = 1 \text{ nM}$, $\mu_I = 0.125 \text{ nM min}^{-1}$, $n = \{1, 10, 100\}$, $\beta_I = \{0.5, 5, 50\} \text{ nM}^{-1} \text{ min}^{-1}$. |
| **Fig. S6B** | FAD v1 (Section S4.2 & Section S2.2) | $\gamma = 0.01 \text{ min}^{-1}$, $\gamma_U = 0.05 \text{ min}^{-1}$, $\gamma_W = 1 \times 10^{-4} \text{ min}^{-1}$, $\eta_0 = 1 \times 10^{-4} \text{ min}^{-1}$, $\eta_- = 0.5 \text{ min}^{-1}$, $\gamma_Y = 1 \text{ min}^{-1}$; and specifically for the double-negative subsystem: $K_D = 1 \text{ nM}$, $\mu_0 = 1.25 \text{ min}^{-1}$, $\mu_1 = 12.5 \text{ nM min}^{-1}$, $K_1 = 1 \text{ nM}$, and for the subsystem with positive feedback: $\mu_0 = 1.25 \text{ min}^{-1}$, $\mu_1 = 12.5 \text{ nM min}^{-1}$, $\mu_P = 10 \text{ nM min}^{-1}$, $K_P = 1 \text{ nM}$; initial parameter values: $\mu_U = 0.125 \text{ min}^{-1}$, $\mu_W = 0.1 \text{ nM min}^{-1}$, $\eta_+ = 0.0375 \text{ nM}^{-1} \text{ min}^{-1}$. |

Note: Across this document, capital letters (e.g. $X$ ) represent both the species and its concentration; the sub-index $X_{ss}$ refers to the steady state value; and lower-case Greek letters represent parameters, which by default are non-negative real numbers.

## S1 CoRa approach

### CoRa

**CoRa** –or ***Co****ntrol* ***Ra****tio*– aims to quantify the effect of feedback control on a system’s ability to reject a step perturbation, while considering the effect and constraints of the individual biochemical events. This is done by directly comparing the feedback system of interest to a locally analogous system without feedback under the formalism of *mathematically controlled comparisons* [1]. Each locally analogous system has exactly the same biochemical reactions and parameters as the original feedback system (i.e. *internal equivalence*), with the exception of the feedback link from the controlled subsystem. For each specific parameter set Θ (i.e. the value of all parameters describing the system of interest), the feedback link is substituted by an equivalent constant input calibrated such that the steady-state of all common species between the two systems are identical before a perturbation is applied (i.e. *external equivalence*). This equivalence allows for a direct comparison of the output change of both systems following a specific step perturbation (e.g. step change in a parameter value), while accounting for the influence of the nonlinearity, saturation, and other intrinsic particularities of the system, and guarantying that any differential response of these two analogous systems represents an *inherent functional difference* associated with the feedback control. The perturbation considered must not affect the constant input of the locally analogous system, as otherwise the differential output response can no longer be uniquely associated with the feedback control.

Let *Y*_*ss*_|_Θ_ denote the steady-state value of the system with feedback for a parameter set Θ, and *Y*_*ss,NF*_ | _Θ_ denote the steady-state value of the locally analogous system without feedback. Let’s also consider a a small step perturbation of a specific parameter *ρ* ∈ Θ (*ρ* → *ρ*′). Following this perturbation, *Y*_*ss*_|_Θ,*ρ*→*ρ*′_ and *Y*_*ss,NF*|Θ,*ρ*→*ρ*′_ denote that new steady-states of the feedback system and locally analogous system without feedback, respectively.

CoRa is then defined as:

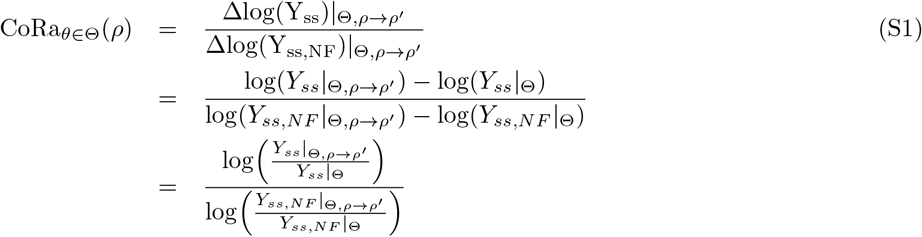

Note that by construction the output of the feedback system and the locally analogous system without feedback are identical before a perturbation, i.e. *Y*_*ss*_|_Θ_ = *Y*_*ss,NF*_ | _Θ_.

Assuming that Δ*ρ* = *ρ*′ − *ρ* is small enough, the output of the feedback system and the locally analogous system without feedback can be expressed as linear functions of Δ*ρ*. The corresponding CoRa function can then be written as:

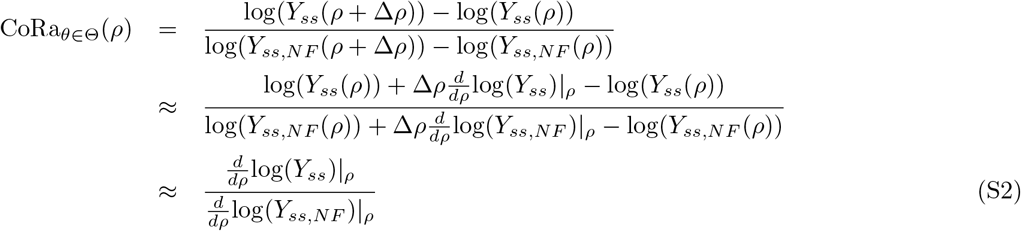

Eq. S2 shows that in this regime, CoRa value is approximately independent of the perturbation size Δ*ρ*. In all the analyses presented on this paper, we used *ρ*′ = 1.05*ρ*. We corroborated that this perturbation size was small enough to reach the linear regime by confirming that identical results were obtained with *ρ*′ = 1.01*ρ*. Nevertheless, with the smaller perturbation size (*ρ*′ = 1.01*ρ*), noise in the numerical solutions was observed for some cases. In general, like for any linearization exercise, the acceptable perturbation size for numerical solutions needs to be evaluated for the specific system and conditions of interest.

The value of CoRa_*θ*∈Θ_(*ρ*) can be easily related to the logic of the feedback (Fig. S1). If CoRa_*θ*∈Θ_(*ρ*) ∈ [0,1), the presence of the feedback reduces the effect of the perturbation compared to the locally analogous system without feedback, i.e. the system has an active negative feedback: either 0 ≤ Δlog(*Y*_*ss*_)|_Θ,*ρ*→*ρ*′_ < Δlog(*Y*_*ss,NF*_)|_Θ,*ρ*→*ρ*′_ or 0 ≥ Δlog(*Y*_*ss*_)|_Θ,*ρ*→*ρ*′_ > Δlog(*Y*_*ss,NF*_) | _Θ,*ρ*→*ρ*′_. On the other hand, if CoRa_*θ*∈Θ_(*ρ*) > 1, the presence of the feedback amplifies the effect of the perturbation compared to the locally analogous system without feedback, i.e. the system has an active positive feedback: either Δlog(*Y*_*ss*_)|_Θ,*ρ*→*ρ*′_ > Δlog(*Y*_*ss,NF*_)|_Θ,*ρ*→*ρ*′_ > 0 or Δlog(*Y*_*ss*_)|_Θ,*ρ*→*ρ*′_ < Δlog(*Y*_*ss,NF*_)|_Θ,*ρ*→*ρ*′_ < 0. Finally, if CoRa_*θ*∈Θ_(*ρ*) = 1, the feedback is effectively inactive. As the goal of CoRa is to quantify feedback control, which by definition requires a corrective (negative) feedback regulation, CoRa_*θ*∈Θ_(*ρ*) is bounded between 0 and 1 for the cases of interest. More specifically, CoRa_*θ*∈Θ_ (*ρ*) = 0 only if the system displays perfect control (*Y*_*ss*_|_Θ,*ρ*→*ρ*′_ = *Y*_*ss*_|_Θ_), and CoRa_*θ*∈Θ_ (*ρ*) value increases as the control effect decreases up until CoRa_*θ*∈Θ_(*ρ*) = 1, when the feedback contribution is effectively zero (i.e. the system response to the perturbation is exactly the same that the one of the system without feedback).

### S1.1 Steps for CoRa implementation

1. Define a solvable **set of ordinary differential equations** representing the biological system of interest, where each equation describes the dynamics of the concentration of a molecular species involved in the system. For example (see Section S2 for the full description of the biological system associated with these equations):

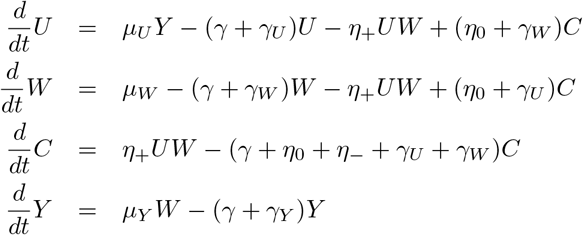
2. Define the **output** of interest, representing a measurement of the controlled subsystem, over which to evaluate the effect of the feedback control; the analysis can be repeated for diverse outputs. In the differential equations example above, and output of interest can be *Y*, and we may be particularly interested in its steady-state value.
3. Determine the **input functions** as all functions dependent on the output defined above (e.g. *Y*) through which this output influences the other molecular species (e.g. *U, W, C*). Input functions are therefore the links from the defined controlled subsystem to the rest of the system. In the example described above, the unique input function (*f*_*θ*_(*Y*)) is the regulated synthesis function of *U* dependent on *Y* :

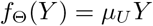
4. Build the **locally analogous no-feedback system** as an identical set of equations as the full, feedback-controlled system, except that the input functions are substituted by constant inputs. These inputs are not dependent on the output (e.g. *Y*), but have identical magnitudes when evaluated in the pre-perturbation steady-state for the given condition (i.e. Θ). We can accomplish this through two alternative strategies:
  a. Introduce some auxiliary species with constitutive expression (i.e. not regulated by any other molecule in our system) with a pre-perturbation steady-state concentration that matches the concentration of the regulatory species in the input functions. Then, use the auxiliary species in the input functions. Using this strategy, the locally analogous no-feedback system for the example described above would be:

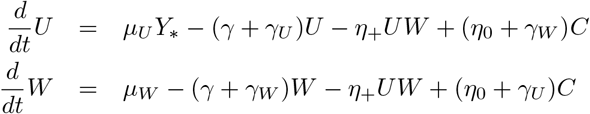

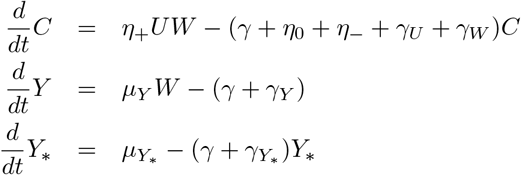

where *Y*_*_ represents the auxiliary (non-regulated) species, which is constitutively expressed with synthesis *μ*_*Y* *_ = *μ*_*Y*_ *W*_*ss*_, and degradation *γ*_*Y* *_ = *γ*_*Y*_, such that the steady-state output of the locally analogous system without feedback *Y*_*ss,NF*_ is equal to the steady state output of the feedback system *Y*_*ss*_ before perturbation to both systems away from that identical steady-state.
  b. Substitute the input functions dependent on the output (e.g. *f*_Θ_(*Y*) = *μ*_*U*_ *Y* in the above example) with a constant whose value is identical to the input function values evaluated at the pre-perturbation steady-state. Using this strategy, the locally analogous no-feedback system for the example described above would be:

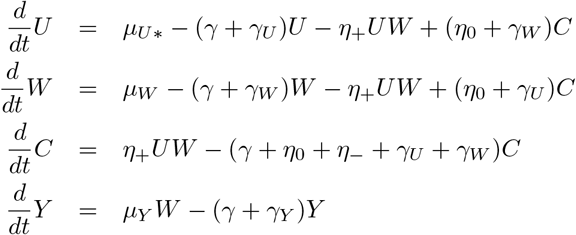

where *U* is now constitutively expressed with synthesis *μ*_*U*\*_ = *μ*_*U*_ *Y*_*ss*_, such that here again, the steady state output of the locally analogous system without feedback *Y*_*ss,NF*_ is equal to the steady state output of the feedback system *Y*_*ss*_. The goal here is that both systems (the original feedback system and its locally analogous system without feedback) have not only identical steady-state values for all species in the condition being evaluated, but if a perturbation occurs, both systems would respond initially in an identical manner, as all the links (regulatory functions) transmit exactly the same information (e.g. with identical levels of non-linearity and saturation), with the clear and intended exception that the “controlled species” in the locally analogous system cannot transmit any feedback information. In general, for this interpretation to be valid, the breaking point (where a regulatory function is substituted by a constant value) must be upstream of where the perturbation occurs; this can be ensured for all types of perturbations if the feedback is broken right where the input function occurs (as proposed here). If the system has multiple feedback loops, and hence multiple input functions need to be defined, we can either evaluate the contribution of each one individually or any combination of them. In either case, the process proceeds exactly as detailed above.
5. Calculate the **output steady-state values** for both systems before and after a perturbation of interest, and obtain the associated CoRa value. For the example described above, for each specific parameter *θ* ∈ Θ:
  a. Calculate the output steady state *Y*_*ss*_|_Θ_ for the original system.
  b. Calculate the output steady state *Y*_*ss,NF*_|_Θ_ for the locally analogous system without feedback system. Confirm that *Y*_*ss*_ = *Y*_*ss,NF*_.
  c. Perturb the desired specific parameter *ρ* ∈ Θ by a small amount, *ρ* → *ρ*′ (e.g. *μ*_*Y*_ → 1.05 *· μ*_*Y*_).
  d. Re-calculate the output steady state *Y*_*ss*_|_Θ,*ρ*→*ρ*′_ for the original system.
  e. Re-calculate the output steady state *Y*_*ss,NF*_|_Θ,*ρ*→*ρ*′_ for the locally analogous system without feedback.
  f. Calculate the CoRa value:

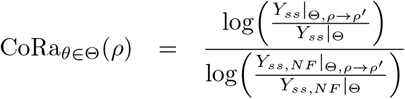
6. Once the system of interest and its locally analogous no-feedback system have been defined, the CoRa analysis can be easily applied over again to **any parameter set and perturbation of interest**:
  a. Update the specific value of the auxiliary species (e.g. *μ*_*Y* *_ = *μ*_*Y*_ *W*_*ss*_) or the constant parameter (e.g. *μ*_*U*\*_ = *μ*_*U*_ *Y*_*ss*_) such that the constant input values in the no-feedback system are identical once again to the input functions values in the feedback system (as described on step #4), keeping the no-feedback system locally-analogous to the feedback system before the perturbation to be evaluated occurs.
  b. Re-calculate the steady-state output responses to the perturbation of interest (step #5).

## S2 Analysis of a modified antithetic feedback control strategy using CoRa

We consider a modified *antithetic feedback motif* (ATF; based on Briat *et al*. [2]) with a simple controlled subsystem consisting of a single molecule *Y*. The ATF motif consists of two molecules *U* and *W* that bind to each other forming a transitory complex *C. C* is then degraded leading to the disappearance of both *U* and *W*. *Y* is produced at a rate that depends on the concentration of *W*, while *U* synthesis is induced by *Y*. The equations of the full system with feedback are then given by:

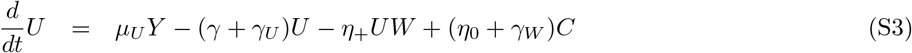

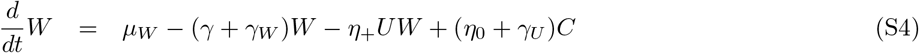

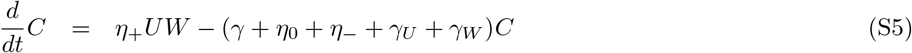

For *Y* dynamics, two alternative scenarios can be easily foreseen: *W* can be either inactivated as a transcription factor once it binds *U* (ATF v1; Fig. 2A,3A),

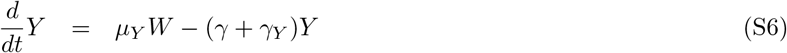

or *W* retains its transcription factor activity until degraded (ATF v2; Fig. 2A,3D),

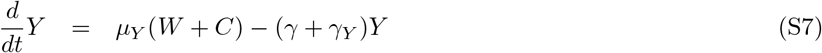

Here all species are subject to loss by dilution (*γ*), in addition of their own individual degradation rates (*γ*_□_), *μ*_□_ represents the synthesis rate for each molecule (either constitutive, *μ*_*W*_, or dependent of a transcription factor, *μ*_*U*_ and *μ*_*Y*_), and *η*_−_ is the co-degradation rate of *U, W* in the complex form *C*; *η*_+_ is the binding rate of *U* and *W* (forming the complex *C*); and *η*_0_ is the spontaneous unbinding rate of these two molecules (dissociating the complex *C*).

Choosing *Y* as the system’s output, the corresponding locally analogous system without feedback maintains the same ODE equations (Eqs. S4-S5, and either Eq. S6 or Eq. S7), with the exception of 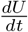,

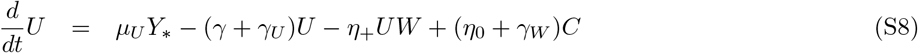

where *U* synthesis rate now depends on a new molecule *Y*_*_ with dynamics

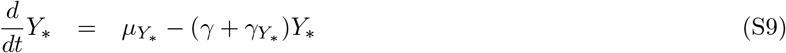

such that *Y*_*_ is constitutively expressed with synthesis *μ*_*Y* *_. If *γ*_*Y* *_ = *γ*_*Y*_, then the steady state output of the locally analogous system without feedback *Y*_*ss,NF*_ is equal to the steady state output of the feedback system *Y*_*ss*_ if either *μ*_*Y* *_ = *μ*_*Y*_ *W*_*ss*_ or *μ*_*Y* *_ = *μ*_*Y*_ (*W*_*ss*_ + *C*_*ss*_), depending on the feedback system being considered (ATF v1 or ATF v2).

In this case, since *Y*_*_ in the locally analogous system without feedback does not depend on any other molecule in the system, its concentration will remain constant after any type of perturbation. As mentioned above, this is an important requirement for the mathematically controlled comparison: if a perturbation also affects *Y*_*_ value (e.g. experimental perturbations on dilution, *γ*), the feedback system and the locally analogous system differ in more than just the feedback information (Fig. S1D), and the CoRa value cannot be interpreted as simply the feedback contribution.

As described by Briat *et al*. [2], assuming there is no dilution (*γ* = 0) as well as no individual degradation of *U* and *W* (i.e. independent of the complex formation *C*; *γ*_*U*_, *γ*_*W*_ = 0), this system (Eqs. S4-S5) is expected to display perfect step disturbance rejection (integral control or perfect adaptation):

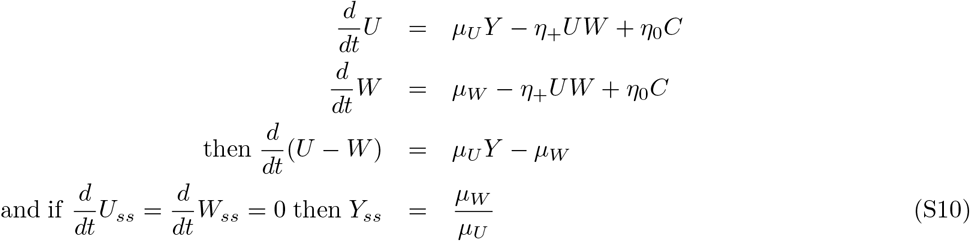

In other words, *Y*_*ss*_ is controlled to a reference value 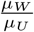, to which it returns exactly after any step perturbation to the system, provided that the steady-state exists and it is stable (see Olsman *et al*. [12] for further discussion). This conclusion is independent of the particular subsystem being controlled, *W* being inactive (Eq. S6) or active (Eq. S7) in the complex form, as well the active degradation rate (*η*_−_), and complex formation dynamics 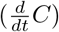.

### S2.1 Understanding effect of saturation on modified antithetic feedback control

#### S2.1.1 ATF control limits with inactive complex

In this section we prove that for the system described in Eqs. S3-S6, if (*γ* + *γ*_*W*_) > 0, as *Y* -synthesis rate (*μ*_*Y*_) value decreases, 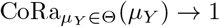. Similarly, if (*γ* + *γ*_*U*_) > 0, as *μ*_*Y*_ increases, CoRa saturates with 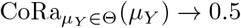. These analytically argued results are corroborated by computational demonstrations in Figure S4A.

##### Proposition 1.

*For the system described in Eqs. S3-S6, as* 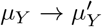, Δlog(*Y*_*ss*_) = Δlog(*μY*) + Δlog(*W*_*ss*_). *Here, for brevity, we denote Y*_*ss*_|_Θ,*μY*_ *by Y*_*ss*_, *and* 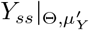 *by* 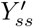, *and similarly for W*_*ss*_. *Therefore* 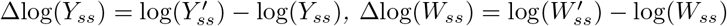, *and* 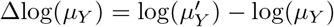.

*Proof*. Given Eq. S6, the output steady state for the system is

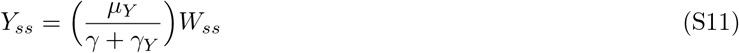

After a perturbation 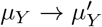, the new output steady state can be written as

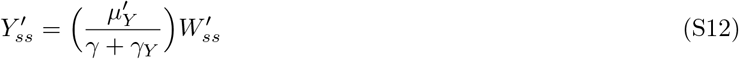

Then, the effect of the perturbation on the system can be quantified as

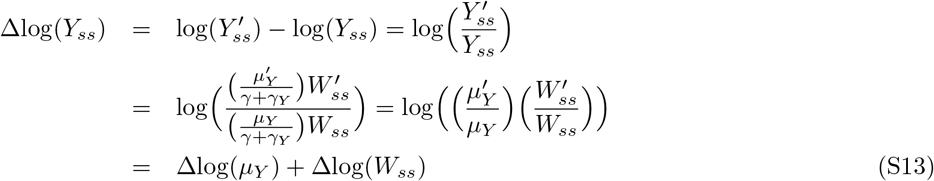

where the effect of the feedback is introduced by the Δlog(*W*_*ss*_) component.

##### Consequence 1.

In the absence of feedback (i.e. when *U* and the *W* do not depend on *Y*), *W*_*ss*_ should remain constant after a *μ*_*Y*_ -perturbation, i.e. Δlog(*W*_*ss*_) = 0. Then, for this system, the effect of the step *μ*_*Y*_ perturbation is simply equal to the size of the perturbation, i.e. Δlog(*Y*_*ss*_) = Δlog(*μ*_*Y*_).

##### Consequence 2.

By definition, a system has feedback control if the presence of feedback reduces the effect of the perturbation over the output change, i.e. |Δlog(*Y*_*ss*_)| < |Δlog(*μ*_*Y*_)|. Then, in order to have feedback control, Δlog(*W*_*ss*_) < 0 if Δlog(*μ*_*Y*_) > 0 (and vice versa). It follows that in a range of *μ*_*Y*_ values with effective feedback control, *W*_*ss*_ must decrease monotonically as *μ*_*Y*_ value increases.

##### Proposition 2.

*For the system described in Eqs. S3-S6, if* (*γ* + *γ*_*W*_) > 0, *the total W steady state (W*_*T,ss*_ = *W*_*ss*_ + *C*_*ss*_*) has an upper limit and lower limit that is independent of μ*_*Y*_. *Additionally, W*_*T,ss*_ *approaches its upper limit when W*_*ss*_ ≈ *W*_*T,ss*_, *and its lower limit when C*_*ss*_ ≈ *W*_*T,ss*_.

*Proof*. Let’s define total *W* as the sum of free molecule *W* and the complex molecule *C*, i.e. *W*_*T*_ = *W* + *C*. Then, the equation of change of *W*_*T*_ corresponds to the sum of Eq. S4 and Eq. S5:

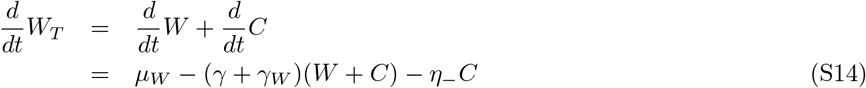

Without loss of generality, we represent *C* as a fraction of the total *W, αW*_*T*_ with *α* ∈ [0, 1]:

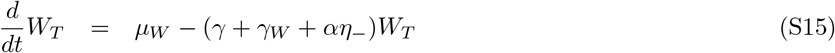

Then, in steady state:

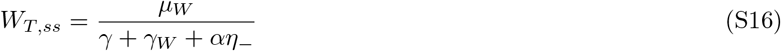

Given that all involved parameters are non-negative, and *α* ∈ [0, 1]:

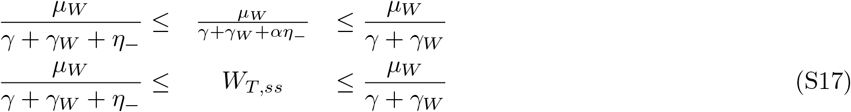

Notice that the upper limit exists only if (*γ* + *γ*_*W*_) > 0. Moreover, it is clear that *W*_*T,ss*_ approaches its upper limit when *α* → 0, i.e. *W*_*T,ss*_ ≈ *W*_*ss*_, while *W*_*T,ss*_ approaches its lower limit when *α* → 1, i.e. *W*_*T,ss*_ ≈ *C*_*ss*_.

##### Proposition 3.

*For the system described in Eqs. S3-S6, and within the range of μ*_*Y*_ *for which the feedback is effective (i*.*e*. |Δlog(*Y*_*ss*_)| < |Δlog(*μ*_*Y*_)| *for all μ*_*Y*_ *values within the range)*, 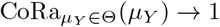 *as μ*_*Y*_ *decreases, provided that* (*γ* + *γ*_*W*_) > 0.

*Proof*. As *W*_*T,ss*_ = *W*_*ss*_ + *C*_*ss*_ is upper bounded (Eq. S17), *W*_*ss*_ must have an upper limit as well (i.e. its supremum, 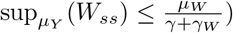. By *Consequence 2* above, within the *μ*_*Y*_ range where feedback control is effective, *W*_*ss*_ value increases as the *μ*_*Y*_ value (before a perturbation is applied) decreases. Therefore, as *μ*_*Y*_ decreases, *W*_*ss*_ approaches its supremum, 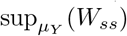. As this occurs, the increment to its concentration (Δlog(*W*_*ss*_)) after an additional perturbation that decreases the *μ*_*Y*_ value even further (i.e. Δlog(*μ*_*Y*_) < 0) is constrained by the *W*_*ss*_ proximity to its limit. With some abuse of notation, we use the symbol ≈ to denote the situation in which this limit is taken as *W*_*ss*_ approaches its upper bound. As a result, in this regime, 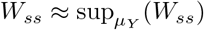 and Δlog(*W*_*ss*_) ≈ 0. Now, using Eq. S13 and *Consequence 1*,

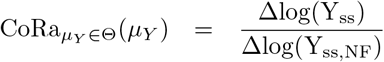

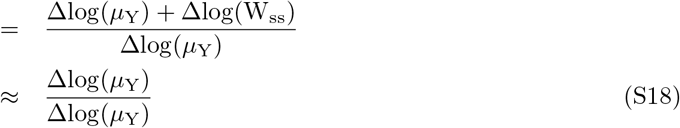

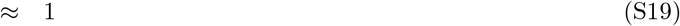

##### Proposition 4.

*For the system described in Eqs. S3-S6, and within the range of μ*_*Y*_ *for which the feedback is effective (i*.*e*. |Δlog(*Y*_*ss*_)| < |Δlog(*μ*_*Y*_)| *for all μ*_*Y*_ *values within the range)*, 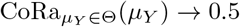 *as μ*_*Y*_ *increases, provided that* (*γ* + *γ*_*U*_) > 0.

*Proof*. By *Consequence 2* above, in a range of *μ*_*Y*_ values with feedback control, *W*_*ss*_ value decreases as the *μ*_*Y*_ value (before a perturbation is applied) increases. As *W*_*T,ss*_ = *W*_*ss*_ + *C*_*ss*_ is lower bounded (Eq. S17), and *W*_*T,ss*_ is minimal when *C*_*ss*_ approaches *W*_*T,ss*_, *C*_*ss*_ must have an lower limit as well (i.e. its infimum, 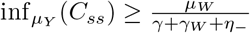), and 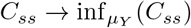 as *μ*_*Y*_ increases.

Let’s define total *U* as the sum of free molecule *U* and the complex molecule *C*, i.e. *U*_*T*_ = *U* + *C*. Then, the equation of change of *U*_*T*_ corresponds to the sum of Eq. S3 and Eq. S5:

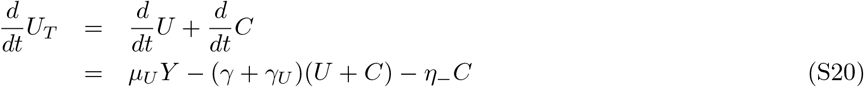

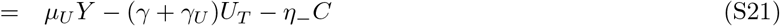

Let’s assume that *μ*_*Y*_ is large enough such that *C*_*ss*_ approaches its lower bound, which is given by 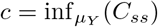. With some abuse of notation, we use the symbol ≈ to denote the situation in which this limit is taken as *C*_*ss*_ approaches its lower bound.

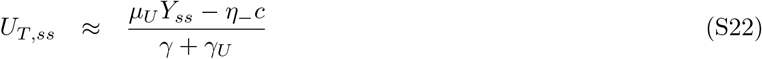

and

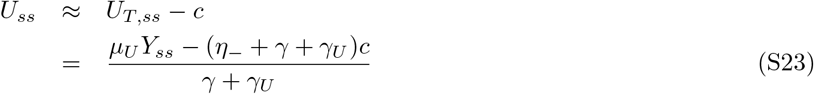

Solving Eq. S5 in steady state, and substituting *C*_*ss*_, *U*_*ss*_,

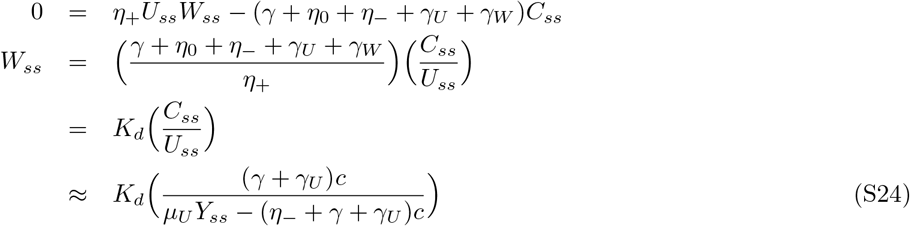

with 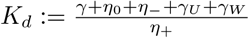. Then, solving Eq. S6 in steady state, and substituting *W*_*ss*_,

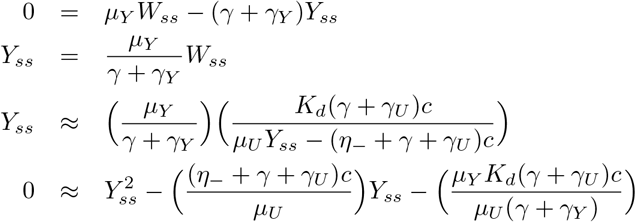

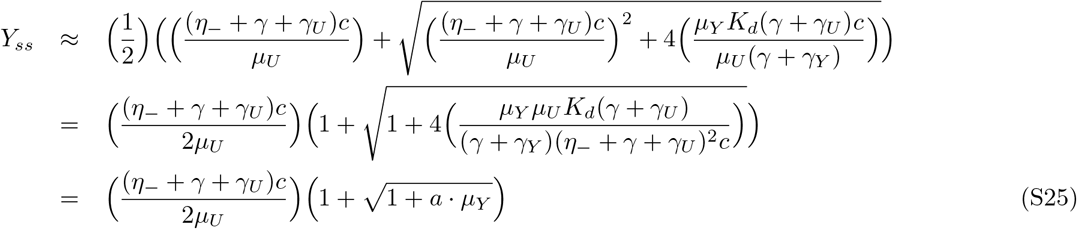

with 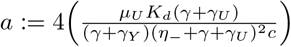. As a result, the change of the steady-state output *Y*_*ss*_ after a small perturbation on *μ*_*Y*_ (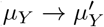, used to compute CoRa),

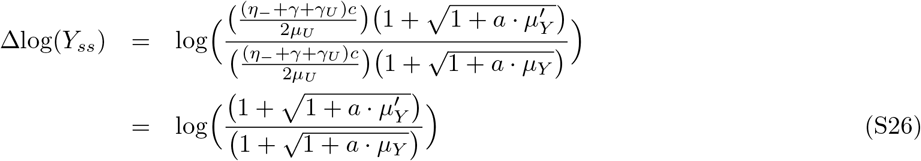

On the other hand, given *Consequence 1*, the no-feedback system has 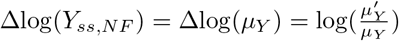, and the associated CoRa value is given by:

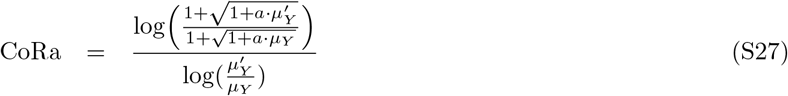

As *μ*_*Y*_ increases, with (*a · μ*_*Y*_) ≫ 1, such that 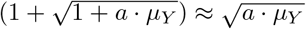, then

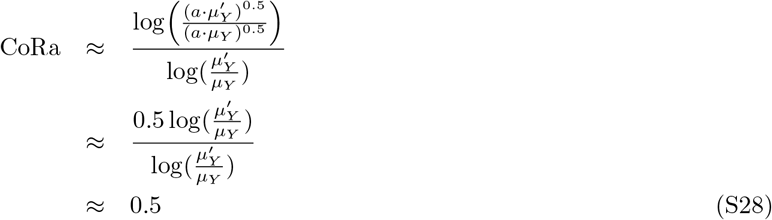

#### S2.1.2 ATF control limits with active complex

In this section, we demonstate that for the system described in Eqs. S3-S5,S7, if (*γ* + *γ*_*W*_) > 0, as *Y* -synthesis rate (*μ*_*Y*_) value decreases, 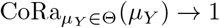. Similarly, as *μ*_*Y*_ increases, CoRa saturates with 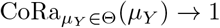, regardless of *γ, γ*_*W*_, *γ*_*U*_ = 0. These analytically argued results are corroborated by computational demonstrations in Figure S4B.

##### Proposition 5.

*For the system described on Eqs. S3-S5,S7, as* 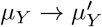, Δlog(*Y*_*ss*_) = Δlog(*μ*_*Y*_) + Δlog(*W*_*T, ss*_). *Here, for brevity, we denote Y*_*ss*_|_Θ,*μY*_ *by Y*_*ss*_, *and* 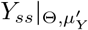 *by* 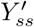, *and similarly for W*_*T,ss*_. *Therefore* 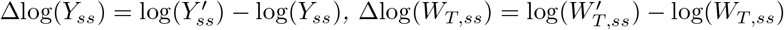, *and* 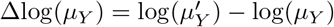.

*Proof*. Given Eq. S7, the output steady state for the system is

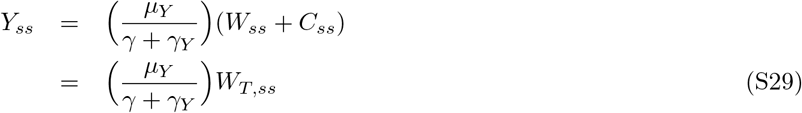

After a perturbation 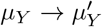, the new output steady state can be written as

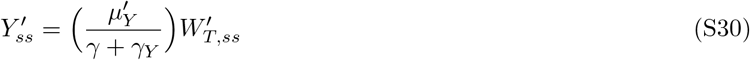

Then, the effect of the perturbation on the system can be quantified as

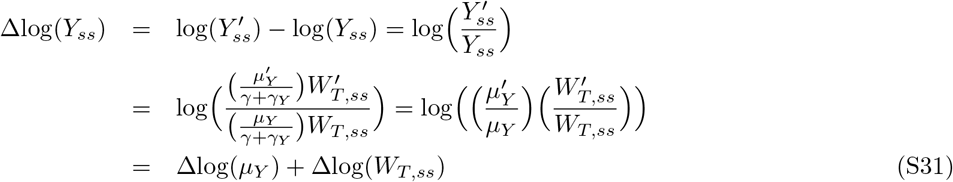

where the effect of the feedback is introduced by the Δlog(*W*_*T,ss*_) component.

##### Consequence 3.

In the absence of feedback (i.e. when *U* and *W* do not depend on *Y*), *W*_*T,ss*_ should remain constant after a *μ*_*Y*_ -perturbation, i.e. Δlog(*W*_*T,ss*_) = 0. As a result, the effect of the perturbation on the system is simply equal to the size of the perturbation, i.e. Δlog(*Y*_*ss*_) = Δlog(*μ*_*Y*_).

##### Consequence 4.

By definition, a system has feedback control if the presence of feedback reduces the effect of the perturbation over the output change, i.e. |Δlog(*Y*_*ss*_)| < |Δlog(*μ*_*Y*_)|. Then, in order to have feedback control, Δlog(*W*_*T,ss*_) < 0 if Δlog(*μ*_*Y*_) > 0 (and vice versa). It follows that in range of *μ*_*Y*_ values with effective feedback control, *W*_*T,ss*_ must decrease monotonically as *μ*_*Y*_ value increases.

##### Proposition 6.

*For the system described in Eqs. S3-S5,S7, if* (*γ* + *γ*_*W*_) > 0, *the total W steady state (W*_*T,ss*_ = *W*_*ss*_ + *C*_*ss*_*) has an upper limit and lower limit, independent of μ*_*Y*_. *Additionally, W*_*T,ss*_ *approaches its upper limit when W*_*ss*_ ≈ *W*_*T,ss*_, *and its lower limit when C*_*ss*_ ≈ *W*_*T,ss*_.

*Proof*. Let’s define total *W* as the sum of free molecule *W* and the complex molecule *C*, i.e. *W*_*T*_ = *W* + *C*. Then, the equation of change of *W*_*T*_ corresponds to the sum of Eq. S4 and Eq. S5:

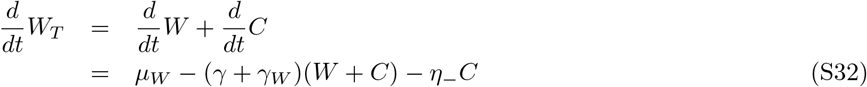

Without loss of generality, we represent *C* as a fraction of the total *W, αW*_*T*_ with *α* ∈ [0, 1]:

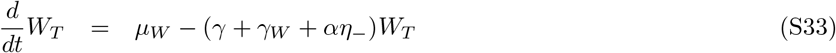

Then, at steady state:

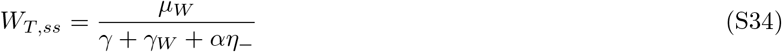

Given that all involved parameters are non-negative, and *α* ∈ [0, 1]:

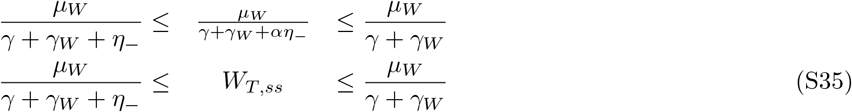

Notice that the upper limit exists only if (*γ* + *γ*_*W*_) > 0. Moreover, it is clear that *W*_*T,ss*_ approaches its upper limit when *α* → 0, i.e. *W*_*T,ss*_ ≈ *W*_*ss*_, while *W*_*T,ss*_ approaches its lower limit when *α* → 1, i.e. *W*_*T,ss*_ ≈ *C*_*ss*_.

##### Proposition 7.

*For the system described in Eqs. S3-S5,S7 and within the range of μ*_*Y*_ *for which the feedback is effective (i*.*e*. |Δlog(*Y*_*ss*_)| < |Δlog(*μ*_*Y*_)|), 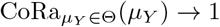 *as μ*_*Y*_ *decreases, provided that* (*γ* + *γ*_*W*_) > 0.

*Proof*. By *Consequence 4*, in the range of effective feedback control, *W*_*T,ss*_ value increases as the *μ*_*Y*_ value (before a perturbation is applied) decreases. Therefore, as the *μ*_*Y*_ value decreases, *W*_*T,ss*_ approaches its limit, 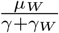 (Eq. S35). Therefore, the potential increment to its concentration (Δlog(*W*_*T,ss*_)) after a perturbation that decreases *μ*_*Y*_ value even further (i.e. Δlog(*μ*_*Y*_) < 0) is constrained by the *W*_*T,ss*_ proximity to the limit. With some abuse of notation, we use the symbol ≈ to denote the situation in which the limit is taken as *W*_*ss*_ approaches its upper bound. In this regime, 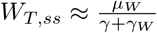 and Δlog(*W*_*ss*_) ≈ 0. Using Eq. S31 and *Consequence 3*,

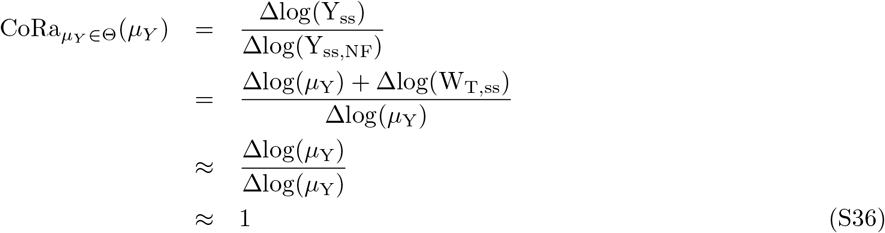

##### Proposition 8.

*For the system described in Eqs. S3-S5,S7, and within a range in which the feedback is effective (i*.*e*. |Δlog(*Y*_*ss*_)| < |Δlog(*μ*_*Y*_)| *for all μ*_*Y*_ *values within the range)*,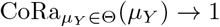 *as μ*_*Y*_ *increases*.

*Proof*. By *Consequence 4* above, in a range of *μ*_*Y*_ values with effective feedback control, *W*_*T,ss*_ value decreases as the *μ*_*Y*_ value (before a perturbation is applied) increases. Therefore, as the *μ*_*Y*_ value increases, *W*_*T,ss*_ approaches its limit, 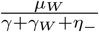 (Eq. S35). Then the potential reduction on its concentration (Δlog(*W*_*T,ss*_)) after a perturbation that increases *μ*_*Y*_ value even further (i.e. Δlog(*μ*_*Y*_) > 0) is constrained by the *W*_*T,ss*_ proximity to the limit. Then as the *μ*_*Y*_ value (before a perturbation is applied) increases, such that 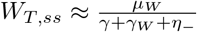 and Δlog(*W*_*ss*_) ≈ 0 (with the same abuse of notation highlighted above as to limits), using Eq. S31 and *Consequence 3*,

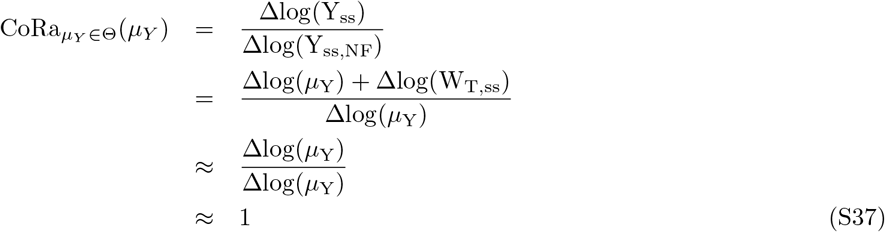

Notice this limit exists even if *W* and *U* are lost only through their mutual annihilation (i.e. *γ, γ*_*W*_, *γ*_*U*_ = 0), as the active degradation is not spontaneous (i.e. 0 < *η*_−_ < ∞).

### S2.2 Limits and the controlled system

It must be emphasized that the control limits described above depend directly on the specific subsystem being controlled, and that analytical intuitive expressions might not always be feasible. CoRa has the advantage of not having to rely on this knowledge. In this paper, we also analyze three different controlled subsystems with the antithetic feedback control (ATF), for which no clear analytical derivations are possible :

1. One-step subsystem:

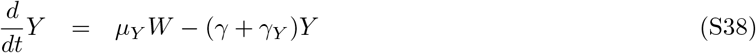
2. Double-negative subsystem:

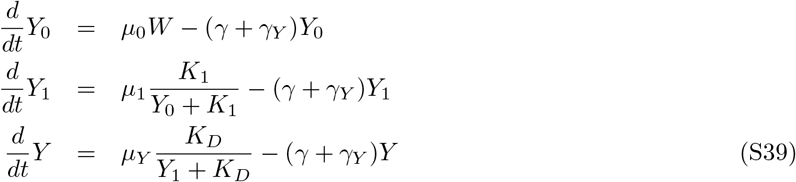
3. Subsystem with positive feedback:

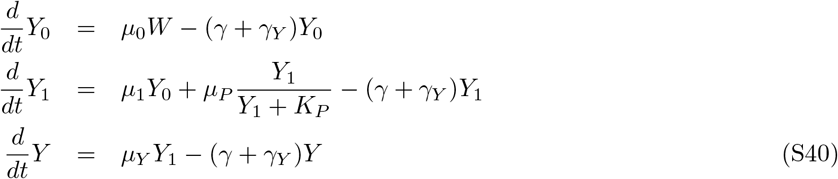

In all cases, *W* induces the synthesis of the subsystem, and *Y* is the output of interest (Eqs. S3-S5; Fig. S2). Choosing *Y* as the system’s output, the corresponding locally analogous system without feedback maintains the same ODE equations except for the input to the control subsystem (*U* synthesis induction for the ATF examples), where *Y* is substituted by a new molecule *Y*_*_, which is constitutively expressed such that the steady state output of the locally analogous system without feedback *Y*_*ss,NF*_ is equal to the steady state output of the feedback system *Y*_*ss*_ (i.e. *Y*_*_ degradation rate *γ*_*Y* *_ = *γ*_*Y*_, and *Y*_*_ synthesis rate, 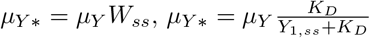, or *μ*_*Y* *_ = *μ*_*Y*_ *Y*_1,*ss*_, depending on the subsystem being considered).

Even with these simple examples, we observed that depending on the subsystem being controlled, the exact same control motif has not only different performance, but qualitatively different responses to the tuning of the control parameters (Fig. S2).

## S3 Understanding effect of saturation on buffering + negative feedback control strategy

### System proposed in Hancock *et al*. (2017)

Hancock *et al*. (2017) explored a simple model proposed to display perfect adaptation. This system consisted of only two species, one working as a buffer of the other while inhibiting its own synthesis (i.e. negative feedback). The equations of this control strategy with a the simple controlled subsystem used in this paper are:

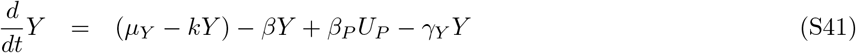

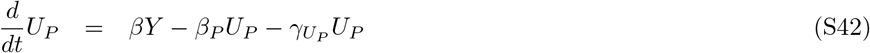

where *μ*_*Y*_ is the maximum synthesis rate of *Y, β* and *β*_*P*_ are inactivation and activation rates respectively, *U*_*P*_ represents the inactive form of *Y, γ*_*Y*_ and 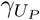 are the degradation rates of *Y* and *U*_*P*_, respectively, and *k* is inhibition rate of *Y* over its own synthesis.

At steady state,

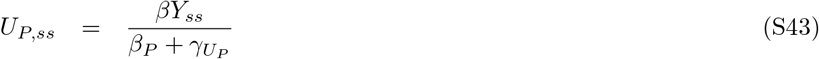

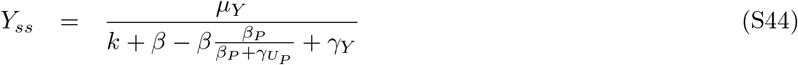

Then, assuming 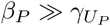, *Y*_*ss*_ is controlled with a reference value 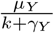.

We consider a modified implementation of this buffering + negative feedback (BNF v1) control motif where the feedback has an additional intermediate step:

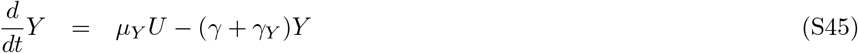

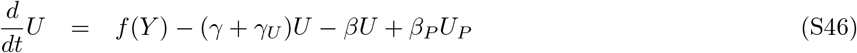

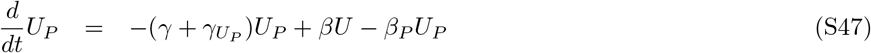

The steady state solution for *U* and *U*_*P*_ is:

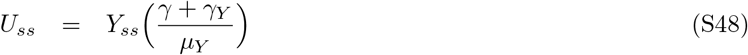

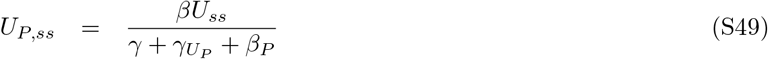

For *Y*, in the case where *f* (*Y*) = *μ*_*U*_ − *kY* is a linear function:

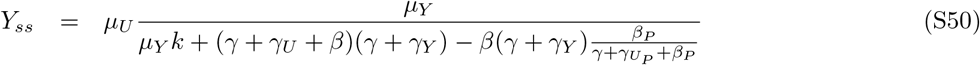

If we assume that 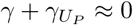, then Eq. S50 is reduced to:

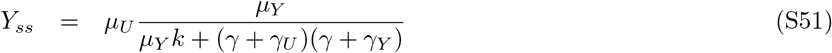

The system has perfect adaptation only if *μ*_*Y*_ *k* ≫ (*γ* + *γ*_*U*_)(*γ* + *γ*_*Y*_), in which case the reference value is 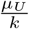.

In the case where 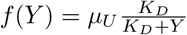 is a Michaelis-Menten function, steady state solution for *Y* is:

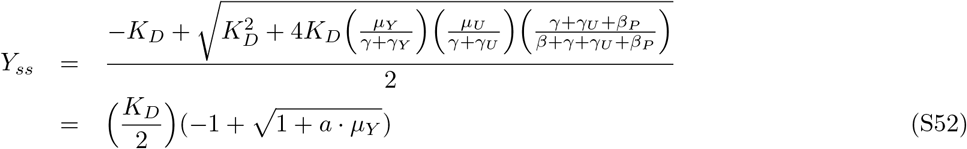

with 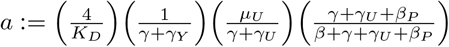. This steady state expression already suggests that perturbations to *μ*_*Y*_ cannot be perfectly controlled anymore. Moreover, we show below that regardless of the parameter values, BNF v1 with a Michaelis-Menten function describing the negative regulation has 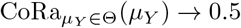.

The corresponding locally analogous system without feedback maintains the same ODE equations (Eq. S45 and Eq. S47), with the exception of 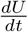,

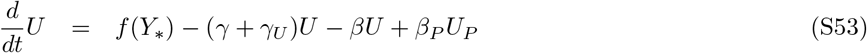

where *U* synthesis rate now depends on a new molecule *Y*_*_ with dynamics

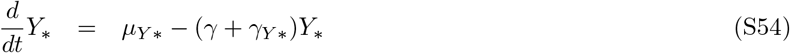

such that, for each parameter set Θ, *Y*_*_ is constitutively expressed with synthesis *μ*_*Y* *_ equal to *Y* synthesis rate in the pre-perturbation steady state solution (i.e. *μ*_*Y* *_ = *μ*_*Y*_ *U*), and degradation rate *γ*_*Y* *_ = *γ*_*Y*_. Then, with 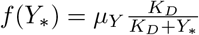, the output steady state solution *Y*_*ss,NF*_ for this locally analogous system without feedback is:

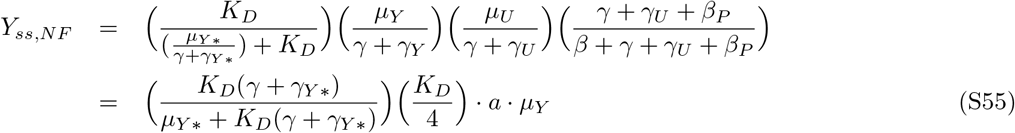

### Control limits

Using Eq. S52 and Eq. S55, the CoRa value for a small perturbation on 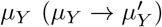 is calculated as,

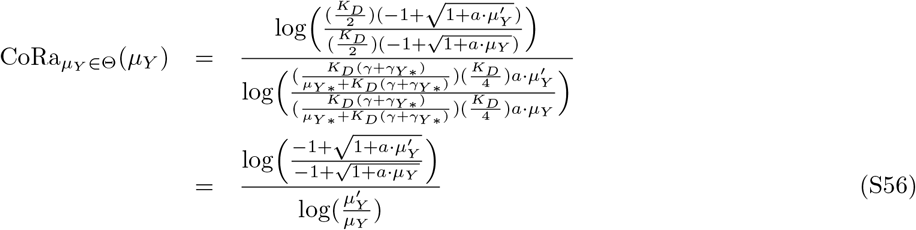

First, we show that 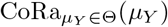 decreases monotonically as the *μ*_*Y*_ value (before the perturbation) increases (i.e. 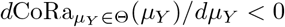). In order to evaluate the derivative of CoRa, we first need to derive the continuous form of the CoRa function (CoRa^*C*^), which corresponds to CoRa evaluated in the limit as the perturbation size (Δ*μ*_*Y*_, with 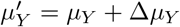) approaches zero,

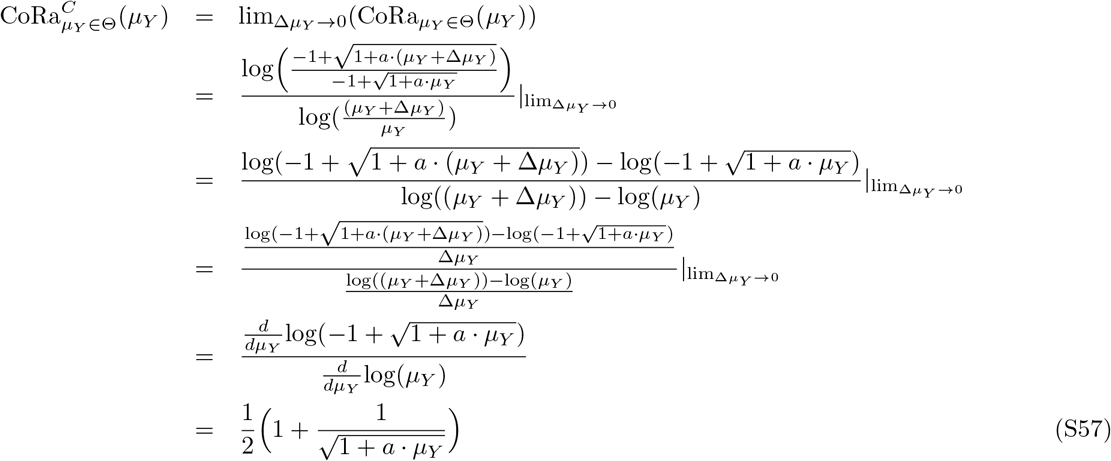

Then,

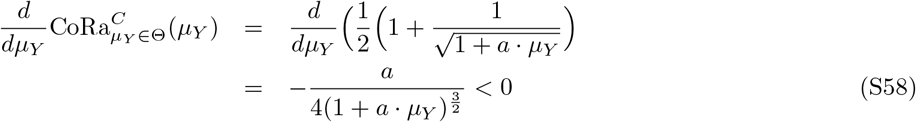

As all parameters are positive (i.e. *a* > 0 and *μ*_*Y*_ > 0), this derivative is always negative.

From Eq. S56, it is easy to see that as the *μ*_*Y*_ value (before the perturbation) increases, with (*a* · *μ*_*Y*_) ≫ 1, such that 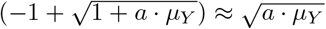, then

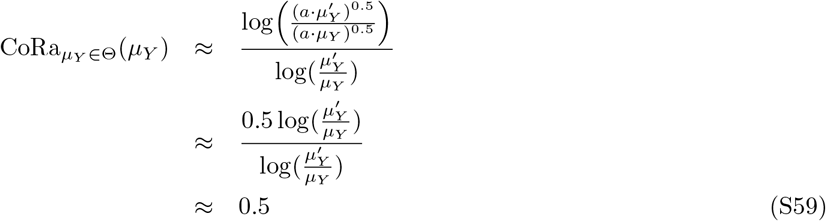

It follows that regardless of the parameter values, BNF v1 with a Michaelis-Menten function describing the negative synthesis regulation has 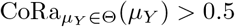.

## S4 Comparing Feedback Control Morifs with CoRa

For all systems below, *Y* represents the system output.

### S4.1 Antithetic Feedback

We consider a simple version of the Antithetic Feedback motif (ATF) proposed by Briat *et al*. [2], where *Y* is being produced at a rate that depends on the concentration of *W*, while *U* synthesis is induced by *Y*, which then binds *W*, forming a transitory complex *C*, which eventually leads to the mutual degradation of *U* and *W* :

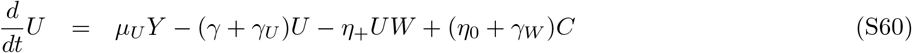

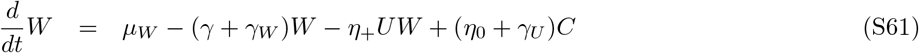

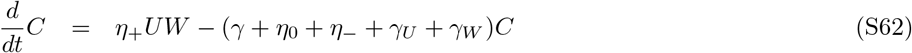

For *Y* dynamics, two alternative scenarios can be easily foreseen: *W* can be either inactivated as a transcription factor once it binds *U* (ATF v1; Fig. 3A),

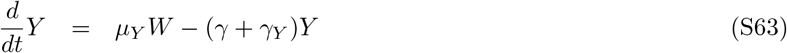

or *W* retains its transcription factor activity until degraded (ATF v2; Fig. 3D),

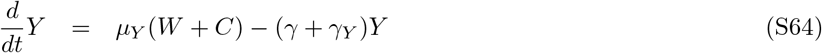

Here all species are subject to loss by dilution (*γ*), in addition of their own individual degradation rates (*γ*_□_), *μ*_□_ represents the synthesis rate for each molecule (either constitutive, *μ*_*W*_, or dependent of the associated transcription factor, *μ*_*U*_ and *μ*_*Y*_), and *η*_−_ is the co-degradation rate of *U, W* in the complex form *C*; *η*_+_ is the binding rate of *U* and *W* (forming the complex *C*); and *η*_0_ is the spontaneous unbinding rate of these two molecules (dissociating the complex *C*).

The corresponding locally analogous system without feedback maintains the same ODE equations (Eq. S61-S62, and either Eq. S63 or Eq. S64), with the exception of 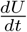,

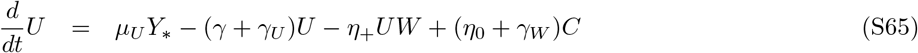

where *U* synthesis rate now depends on a new molecule *Y*_*_ with dynamics

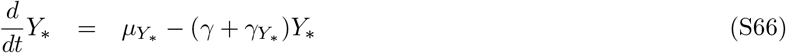

For each parameter set Θ, *Y*_*_ is constitutively expressed with synthesis *μ*_*Y* *_ equal to *Y* synthesis rate in the pre-perturbation steady state solution (i.e. either *μ*_*Y* *_ = *μ*_*Y*_ *W*_*ss*_ or *μ*_*Y* *_ = *μ*_*Y*_ (*W*_*ss*_ + *C*_*ss*_), depending on the feedback system being considered), and degradation rate *γ*_*Y* *_ = *γ*_*Y*_.

### S4.2 Feedback by Active Degradation

We consider a simple version of the Feedback by Active Degradation motif (FAD; [11, 16]), where *Y* is being produced at a rate that depends on the concentration of *W*, while *U* synthesis is induced by *Y*. *Y* then binds *W*, forming a transitory complex *C*, which eventually leads to the degradation of only *W* while freeing *U* :

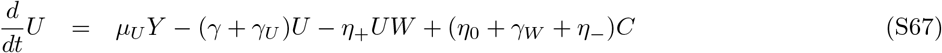

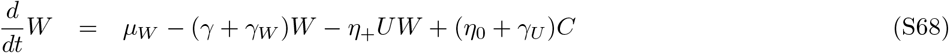

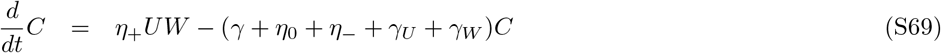

For *Y* dynamics, two alternative scenarios can be easily foreseen: *W* can be either inactivated as a transcription factor once it binds *U* (FAD v1; Fig. 3B),

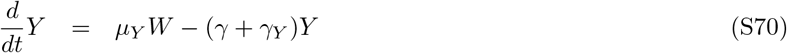

or *W* retains its transcription factor activity until degraded (FAD v2; Fig. 3E),

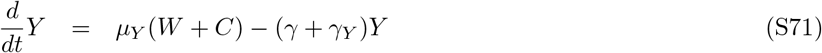

Here all species are subject to loss by dilution (*γ*), in addition of their own individual degradation rates (*γ*_□_), *μ*_□_ represents the synthesis rate for each molecule (either constitutive, *μ*_*W*_, or dependent of the associated transcription factor, *μ*_*U*_ and *μ*_*Y*_), and *η*_−_ is the active degradation rate of *W* in the complex form *C*; *η*_+_ is the binding rate of *U* and *W* (forming the complex *C*); and *η*_0_ is the spontaneous unbinding rate of these two molecules (dissociating the complex *C*).

The corresponding locally analogous system without feedback maintains the same ODE equations (Eq. S68-S69, and either Eq. S70 or Eq. S71), with the exception of 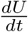,

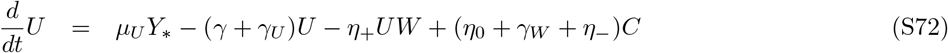

where *U* synthesis rate now depends on a new molecule *Y*_*_ with dynamics

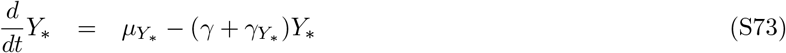

For each parameter set Θ, *Y*_*_ is constitutively expressed with synthesis *μ*_*Y* *_ equal to *Y* synthesis rate in the pre-perturbation steady state solution (i.e. either *μ*_*Y* *_ = *μ*_*Y*_ *W*_*ss*_ or *μ*_*Y* *_ = *μ*_*Y*_ (*W*_*ss*_ + *C*_*ss*_), depending on the feedback system being considered), and degradation rate *γ*_*Y* *_ = *γ*_*Y*_.

### S4.3 Feedback by Active Degradation + Positive Feedback with inactive complex

We consider the FAD motif with the addition of a positive feedback (FDP; [4, 16]), i.e. *W* induces its own synthesis. Once again, two alternative scenarios can be easily foreseen: *W* can be either inactivated as a transcription factor once it binds *U* (FDP v1; Fig. 3C),

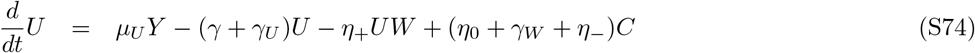

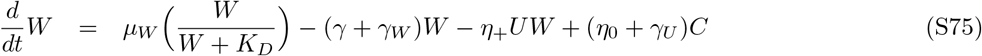

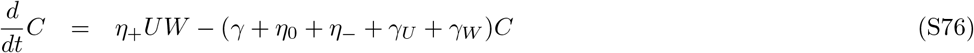

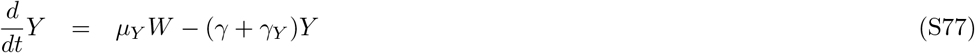

or *W* retains its transcription factor activity until degraded (FDP v2; Fig. 3F),

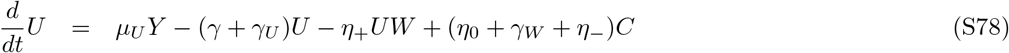

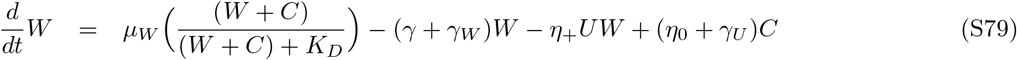

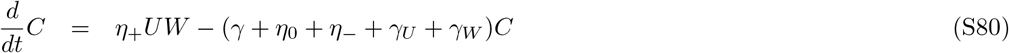

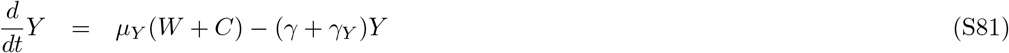

Here all species are subject to loss by dilution (*γ*), in addition of their own individual degradation rates (*γ*_□_), *μ*_□_ represents the synthesis rate for each molecule, *K*_*D*_ is the Michaelis-Menten constant for *W* auto-regulation, and *η*_−_ is the active degradation rate of *W* in the complex form *C*; *η*_+_ is the binding rate of *U* and *W* (forming the complex *C*); and *η*_0_ is the spontaneous unbinding rate of these two molecules (dissociating the complex *C*).

The corresponding locally analogous system without feedback maintains the same ODE equations (either Eq. S75-S77, or Eq. S79-S81), with the exception of 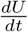,

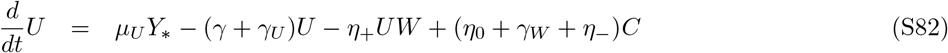

where *U* synthesis rate now depends on a new molecule *Y*_*_ with dynamics

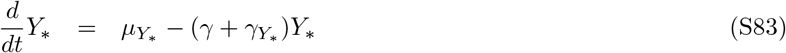

For each parameter set Θ, *Y*_*_ is constitutively expressed with synthesis *μ*_*Y* *_ equal to *Y* synthesis rate in the pre-perturbation steady state solution (i.e. either *μ*_*Y* *_ = *μ*_*Y*_ *W*_*ss*_ or *μ*_*Y* *_ = *μ*_*Y*_ (*W*_*ss*_ + *C*_*ss*_), depending on the feedback system being considered), and degradation rate *γ*_*Y* *_ = *γ*_*Y*_.

### S4.4 Buffering + Negative Feedback

We consider a motif with negative feedback and a buffering loop (BNF v1 & v2; Fig. 3G-H), similar to the one proposed in Hancock *et al*. [6], where *Y* represses the synthesis of *U*, and *U* transitions to an alternative state *U*_*P*_ and vice versa:

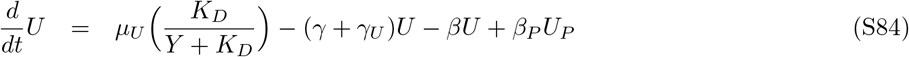

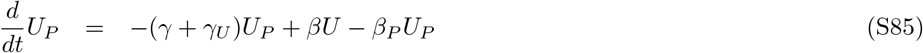

closing the feedback with either *U* inducing *Y* synthesis (BNF v1; Fig. 3G),

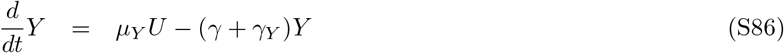

or *U*_*P*_ inducing *Y* synthesis (BNF v2; Fig. 3H):

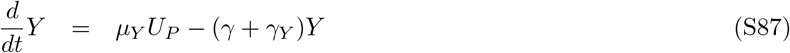

Here all species are subject to loss by dilution (*γ*), in addition of their own individual degradation rates (*γ*_*Y*_ for *Y*, and *γ*_*U*_ for both *U* and *U*_*P*_), *μ*_*U*_ is the maximum synthesis rate of *U* (in absence of *Y*), *μ*_*Y*_ is the synthesis rate of *Y* (depending either on *U*, Eq. S86, or *U*_*P*_, Eq. S87), and *β, β*_*P*_ are the transition rates from *U* to *U*_*P*_, and viceversa.

The corresponding locally analogous system without feedback maintains the same ODE equations (Eq. S85, and either Eq. S86 or Eq. S87), with the exception of 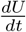,

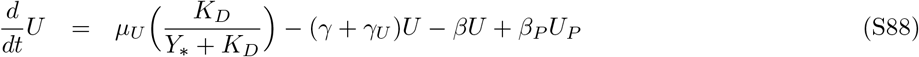

where *U* synthesis rate now depends on a new molecule *Y*_*_ with dynamics

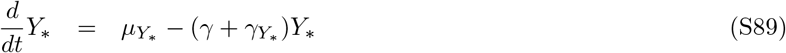

For each parameter set Θ, *Y*_*_ is constitutively expressed with synthesis *μ*_*Y* *_ equal to *Y* synthesis rate in the pre-perturbation steady state solution (i.e. either (i.e. either *μ*_*Y* *_ = *μ*_*Y*_ *U*_*ss*_ or *μ*_*Y* *_ = *μ*_*Y*_ *U*_*P,ss*_, depending on the feedback system being considered), and degradation rate *γ*_*Y* *_ = *γ*_*Y*_.

### S4.5 Feedback + Feedforward Loop

We consider a motif with negative feedback and a coherent feed-forward loop (FFL; Fig. 3H), similar to the one proposed in Harris *et al*. [7], where *Y* represses the synthesis of *U*, and *U* induces the synthesis of both *Y* and *W*, which in turns also induces *Y* synthesis:

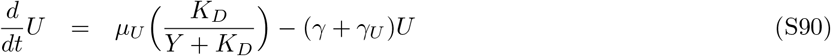

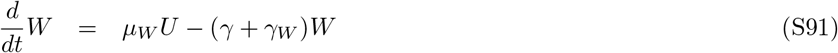

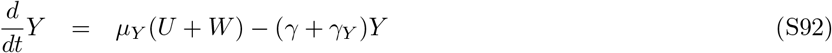

Here all species are subject to loss by dilution (*γ*), in addition of their own individual degradation rates (*γ*_□_), and *μ*_□_ represents the synthesis rate for each molecule.

The corresponding locally analogous system without feedback maintains the same ODE equations (Eq. S91-S92), with the exception of 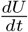,

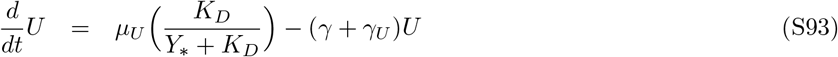

where *U* synthesis rate now depends on a new molecule *Y*_*_ with dynamics

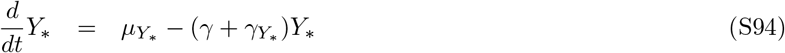

For each parameter set Θ, *Y*_*_ is constitutively expressed with synthesis *μ*_*Y* *_ equal to *Y* synthesis rate in the pre-perturbation steady state solution (i.e. *μ*_*Y* *_ = *μ*_*Y*_ (*U*_*ss*_ + *W*_*ss*_)), and degradation rate *γ*_*Y* *_ = *γ*_*Y*_.

### S4.6 Brink Motif Feedback

We consider a simple version of the Brink motif (BMF) proposed by Samaniego & Franco [14], where *A* and *I* bind and annihilate each other (by creating the complex *C*), *A* induces the activation of *U* (*U*_*P*_ to *U*), while *I* induces its inactivation (*U* to *U*_*P*_), and *U* induces the synthesis of *Y* :

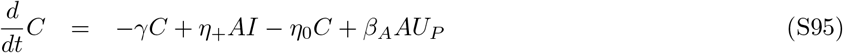

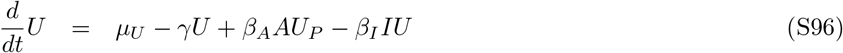

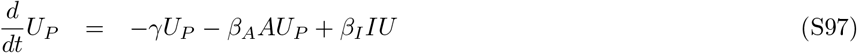

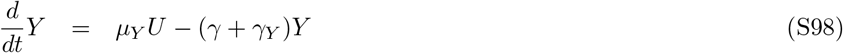

With *Y* either inducing the synthesis of *I* (BMF v1; Fig. 3I),

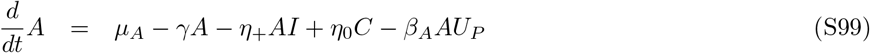

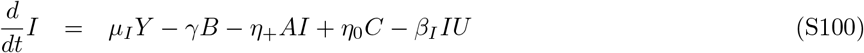

or *Y* repressing the synthesis of *A* (BMF v2; Fig. 3J),

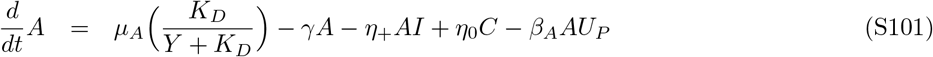

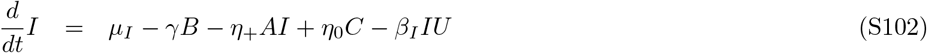

Here all species are subject to loss by dilution (*γ*), *μ*_□_ represents the synthesis rate for each molecule (except *U*_*P*_, which is only created by the inactivation of *U*), *η*_+_ is the binding rate of *A* and *I* (forming the complex *C*), *η*_0_ is the spontaneous unbinding rate of these two molecules (dissociating the complex *C*); and *β*_*A*_, *β*_*I*_ are the activation and inactivation rates of *U*, respectively. Finally, *K*_*D*_ is the Michaelis-Menten constant for the transcriptional repression by *Y* on Eq. S102.

The corresponding locally analogous system without feedback maintains the same ODE equations (Eq. S95-S97, and either Eq. S99 or Eq. S102), with the exception of 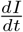 for BMF v1,

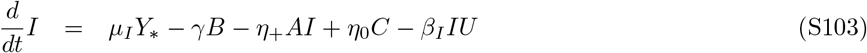

or 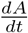 for BMF v2,

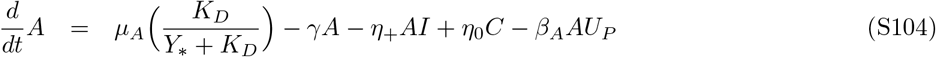

where *I, A* synthesis rate, respectively, now depends on a new molecule *Y*_*_ with dynamics

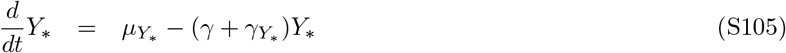

such that *Y*_*_ is constitutively expressed with synthesis *μ*_*Y* *_ equal to *Y* synthesis rate in the steady state solution for each parameter set Θ (i.e. either *μ*_*Y* *_ = *μ*_*Y*_ *U*_*ss*_), and degradation rate *γ*_*Y* *_ = *γ*_*Y*_, before the perturbation.

## S5 Using CoRa to design biomolecular feedback control mechanisms

Below, we present the details of the optimization of the CoRa function over control parameters. For this, we implemented a simple algorithm with two optimization phases: choosing for parameter values that (1) reduce the CoRa value up until min(CoRa) ≤ *ϵ* (with *ϵ* being a threshold picked by the user), and then (2) expand the range of parameter set values *θ* (e.g. range of *μ*_*Y*_ values) with min(CoRa) ≤ *ϵ*. Multiple parameter sets might result in equivalent efficient control for a given feedback control system. This can be explored computationally by running the optimization algorithm for multiple initial conditions and/or random number chains. Iterations of the optimization process allow to determine the region of the parameter space and relationship between parameters associated to the optimal performance for the case of interest.

### S5.1 Optimizing feedback control designs

The goal is to maximize the range of values of a specific parameter *θ* ∈ Θ where CoRa_*θ*∈Θ_(*ρ*) ≤ *ϵ*. For this, we consider two phases of the optimization: first minimizing the min(CoRa_*θ*∈Θ_(*ρ*)) up until it is less or equal *ϵ*; then maximizing the magnitude of | CoRa_*θ*∈Θ_(*ρ*) ≤ *ϵ* | in the explored range (in the logarithmic scale).

#### Error function, *χ*^2^

##### Minimizing min(CoRa_*θ*∈Θ_(*ρ*))

We define our error function (sum of square errors) by assuming the optimal point *D* = 0, and considering the expected variance of uniform distribution ∼ *U* [0, 1] (*σ*^2^ = 0.083). Then our error function in the initial phase of the optimization is:

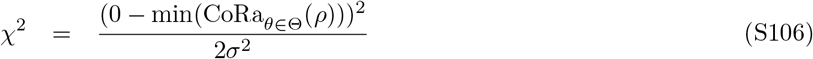

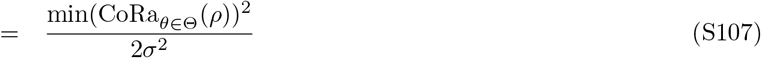

##### Maximizing | CoRa_*θ*∈Θ_(*ρ*) ≤ *ϵ* |

We assume the optimal point *D* = 1 for all values of *θ* ∈ Θ in the range of interest, then

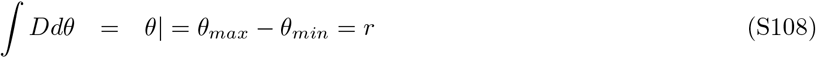

And for each data point *θ*_*i*_, *y*_*i*_ is 1 if 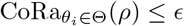, 0 otherwise. Then (*D*_*i*_ − *y*_*i*_)^2^ = 0 for the range where CoRa_*θ*∈Θ_(*ρ*) ≤ *ϵ*, 1 otherwise. Finally,

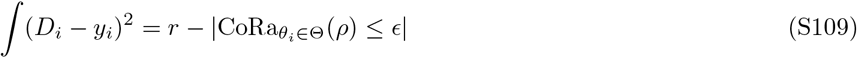

And the range of interest is maximized as this value is minimized. Then, our error function in this phase of the optimization is:

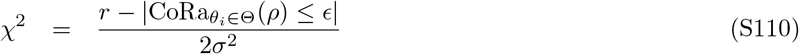

We initially tried using the variance of a uniform function ∼ *U* [0, *r*] (*σ*^2^ = 0.083*r*^2^) for the error function, but it resulted in very noisy simulations. So we opted for the same variance than when minimizing min(CoRa_*θ*∈Θ_(*ρ*))(*σ*^2^ = 0.083).

#### Metropolis Random Walk algorithm

For each phase, an error function is defined, and a Metropolis Random Walk algorithm implemented as follows:

1. Choose some initial parameters Θ_1_ and calculate the corresponding likelihood.
2. Iterate over *t* = *{*1, 2, …, *t*_*MAX*_ *}* as follows:
  a. Draw a random proposal 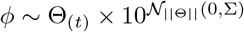 where 𝒩_‖Θ‖_(0, Σ) is a Multivariate Normal distribution with the same dimension as Θ_(*t*)_, mean zero and covariance matrix Σ = 0.1.
  b. We construct a likelihood function using a Gaussian function:

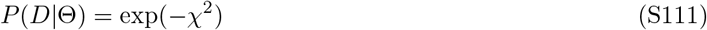

where Θ is the set of parameter to be optimized, *D* is the optimal data, and *χ*^2^ is the error function (which depends on the optimization phase). Note the likelihood is maximal when the error is minimal. Then we calculate the likelihood ratio:

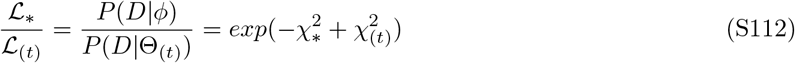 Accept the proposed *ϕ* if the ratio is larger than a random number ∼ *U*[0, 1]. The proposed value is always accepted if the error is smaller (i.e. it’s better).
  c. Update parameters Θ_(*t*+1)_ ← *ϕ* with probability 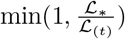; otherwise, Θ_(*t*+1)_ ← Θ_(*t*)_.

